# Investigating the barley methylome, its variation and association with genomic, transcriptomic, and phenotypic variation

**DOI:** 10.1101/2024.10.21.619366

**Authors:** Marius Kühl, Po-Ya Wu, Asis Shrestha, Julia Engelhorn, Thomas Hartwig, Benjamin Stich

**Affiliations:** Institute of Quantitative Genetics and Genomics of Plants, Heinrich Heine University, 40225 Düsseldorf, Germany.; Institute for Breeding Research on Agricultural Crops, Julius Kühn Institute, Federal Research Centre for Cultivated Plants, 18190 Sanitz, Germany.; CEPLAS Cluster of Excellence on Plant Sciences, 40225 Düsseldorf, Germany.; Max Plank Institute for Plant Breeding Research, 50829 Cologne, Germany.

## Abstract

Epigenetic variation contributes to explaining the missing heritability of complex traits. In order to understand the genome-wide methylation variation in spring barley, our objectives were to gain fundamental insight into the barley methylome through whole genome bisulfite sequencing, characterizing methylation variation among 23 parental inbreds of a community resource for genetic mapping of phenotypic traits, and assessing the association of differentially methylated regions (DMRs) with single nucleotide polymorphisms (SNPs) and gene expression variation. Compared to other angiosperms, barley was found to have a highly methylated genome with an average genome wide methylation level of 88.6%, 58.1%, and 1.4% in the CpG, CHG, and CHH sequence context, respectively. We identified just below 500 000 differentially methylated regions (DMRs) among the inbreds. About 64%, 64%, and 83% of the DMRs were not associated with genomic variation in the CpG, CHG, and CHH context, respectively. The methylation level of around 6% of all DMRs was significantly associated with gene expression, where the directionality of the correlation was depended on the relative location of the DMR to the respective gene with a recognizable pattern. Notably, this pattern was much more specific and spatially confined than the association of methylation with gene expression across genes in a singular inbred line. We exemplified this association between DNA methylation and gene expression on the known flowering promoting gene *VRN-H1* and identified a highly methylated epiallele associated with earlier flowering time. Finally, methylation was shown to improve the prediction abilities of genomic prediction models for a variety of traits over models using solely SNPs and gene expression as predictors. These observations highlight the independence of DNA methylation to sequence variation and their difference in information content. Our discoveries suggest that epigenetic variation provides a layer of information likely not predictable by other means and is therefore a valuable addition to genomic prediction models.

## INTRODUCTION

Barley (*Hordeum vulgare*) is one of the world’s oldest and most produced crops with worldwide importance for animal feed and alcohol production [1]. In addition, it is not only important for global food security, but also serves as an important model organism for other cereals due to its smaller diploid genome and simple inbreeding genetics [2]. This raises the need for a greater understanding of its genetics, genomics, and physiology to increase the yield and meet global demands. However, in any species, genetic variation alone cannot explain all of the phenotypic variation of any macroscopic or microscopic phenotype, a phenomenon known as missing heritability [3]. Epigenetic variation was proposed as one of the possible molecular reasons for it [4].

Many definitions of epigenetics have been proposed over the years [5, 6]. However, today it is commonly accepted that epigenetics summarizes those variations that are not necessarily associated with changes in the DNA sequence, which includes chromatin and histone modifications, non-coding RNAs, and DNA methylation [5, 7–9]. Histone modifications are involved in many important processes like plant development and stress response by primarily recruiting chromatin remodeling enzymes, which effects the accessibility of chromatin and, thus, regulating gene expression [10, 11]. Like histone modifications, noncoding RNAs are involved in many important processes including plant growth, development, stress response, cell differentiation, and cell cycle progression through transcriptional and post-transcriptional gene expression regulation [11, 12]. Both histone modifications and non-coding RNAs regulate the response to biotic and abiotic stresses in barley and its close relative wheat [11]. For example, during drought stress, barley increases its H3-density at response genes and establishes euchromatic marks that increase their expression [13].

One of the most researched aspects of epigenetics is DNA methylation. This is because evidence for its heritability is rather well-established in comparison to other epigenetic variations [7, 14–16]. DNA methylation is the addition of a methyl or hydroxymethyl group to the C5 position of cytosine forming 5-methylcytosine or 5-hydroxymethylcytosine, respectively [17]. In plants, methylation occurs in three sequence context: CpG, CHG (where H refers to any base but G), and CHH [18]. De novo methylation of cytosines is catalyzed by DOMAINS REARRANGED METHYLTRANSFERASE 2 (DRM2) [19]. CpG methylation is then maintained by DNA METHYLTRANSFERASE 1, CHG by CHROMOMETHYLASE, and the asymmetric CHH methylation is maintained by continuous de novo methylation by DRM2. DNA methylation in plants largely targets repetitive elements like transposons [19–21]. Genes are usually only CpG methylated except for transcription start sites (TSS) and transcription termination sites (TTS) where methylation is absent.

Investigation of the methylation landscape is important due to its various regulatory functions. It plays an important role in plant development through zygotic gene expression regulation and imprinting, resulting in the differential expression of maternal or paternal alleles [22, 23]. It is a conserved epigenetic modification that can alter the state of chromatin and therefore influence chromosome interactions. For example, hypermethylation decreases the frequency of crossovers [24]. Methylation can silence transposable elements (TEs) and consequently contribute to the overall genome stability [25]. Furthermore, promoter methylation can lead to downregulation of gene expression by inhibiting transcription activators, promoting repressors or even influencing histone modifications. On the other hand, gene body methylation can increase gene expression, which is strongly conserved between orthologs and therefore a long-term property of evolutionary consequence [21, 26]. For example, DNA methylation regulates many abiotic stress responses in plants and is able to produce a shortand long-term memory without sequence changes [27] and, thus, such epialleles may be inherited independently from the DNA sequence [28]. Barley has been shown to also regulate its drought and moisture stress responses by DNA methylation [29, 30]. In addition, a study of DNA methylation changes in rice suggested that environmentally induced gene expression changes are able to cause DNA methylation changes in nearby TEs at a later time point [31]. In our study, however, we were interested in environment independent methylation differences as a heritable characteristic of inbred lines.

Finding differences in methylation between individuals is usually done by first testing for differentially methylated cytosines (DMCs) and then collapsing regions of multiple DMCs into differentially methylated regions (DMRs), but DMRs can also be identified directly [32]. Interestingly, about half of the common DMRs in a maize study were not associated with local genetic state [33]. In a second study in maize by Xu *et al.* [34] even 60% of the DMRs showed no association with sequence variation. However, these DMRs were strongly correlated with gene expression and even associated with phenotypic traits that could not be explained by single nucleotide polymorphisms (SNPs). Human genetics show that especially the combination of genetic and epigenetic information is improving the phenotypic prediction score for complex traits [35]. Studies with plants also indicate a huge benefit in the addition of epigenetics to phenotypic prediction for many important traits especially seed quality, yield components, energy-use efficiency, and respiration, which are of high interest to plant breeders [36–39]. For example, in a study aiming to predict plant height of *Arabidopsis thaliana*, epigenetic variation explained 65% of the phenotypic variance [40]. However, such information is not yet available for any small grain cereal.

Several studies of DNA methylation in barley are available. However, most of these studies focused on finding differences among two groups, typically a treatment and a control group. For example, the mediation of heat or drought stress through methylation was frequently assessed in different barley varieties [29, 41–43]. While these studies provide insight into methylation changes in barley upon exposure to stress, they fail to characterize the methylome variation among inbred lines, as well as its association with genomic, trancriptomic, and phenotypic variation on a broad scale.

First efforts have been made by Malinowska *et al.* [44] to characterize methylome variation in barley, however the study was based on reduced representation bisulfite sequencing (RRBS) only covering 0.7% of the barley genome. The same limitation is also true for the study of Hansen *et al.* [45]. Therefore, in order to understand the genome-wide methylation variation and support quantitative trait loci cloning projects that rely on the alleles segregating among the 23 spring barley inbreds, the objectives of this study were to:

i. Gain fundamental insight into the spring barley methylome through whole genome bisulfite sequencing (WGBS)
ii. Characterize its variation among 23 parental inbreds of a community resource for mapping phenotypic traits
iii. Assess the association of DMRs with SNPs and gene expression variation

## MATERIAL AND METHODS

### Plant material

From a total of 224 spring barley accessions [46], 23 were selected for maximal combined phenotypic and genotypic richness as previously described by Weisweiler *et al.* [47].

These inbreds are the parental inbreds of the barley double round robin population (Hv-DRR, [48]). Seeds of the 23 inbreds were sown in a greenhouse in mid-July. Seedlings were placed in the vernalization chamber with 16 hours of light at 22 °C for five weeks and subsequently repotted to ensure homogeneous growth. Tissue samples of 3 × 1 cm were collected from the youngest leaf one week after the transfer to the greenhouse. Apex samples were collected when stage 47 of the Zadoks scale was reached [49]. Seedling samples were grown under the same conditions in petri dishes and collected five days after germination. HOR7985, HOR8160, Sissy, SprattArcher, Unumli-Arpa and W23829/803911 tissue samples were mixed, respectively. The remaining inbred line samples consist only of leaf tissue (Table S1). For Sissy, three additional samples were collected: one for leaf, apex and seedling, respectively. The data set comprised replicates only for one sample mimicking the partial replicated design frequently used in a plant breeding context.

### SNP and gene expression data

A set of 79 348 211 SNPs from whole genome DNA sequencing for the 23 barley inbred lines was available from Weisweiler *et al.* [50]. In order to reduce the covariance due to environmental factors between methylation and gene expression, we used two gene expression datasets that were available from Weisweiler *et al.* [47] which were generated for the same genotypes but different plants. This had the advantage that observed correlations are genotypic correlations as they are not caused by common environmental effects [51]. The first dataset consisted of leaf expression data for 21 inbred lines, not including Kombyne and Sanalta. The second set consisted of seedling expression data for 21 inbred lines, not including IG128216 and Sanalta. Raw data was remapped against the Morex v3 reference and subsequently normalized using DESeg2 [52]. Finally, missing inbreds were mean imputed.

### Phenotypic data

Six phenotypic traits were assessed in seven environments (Cologne, 2017 to 2019; Mechernich and Quedlinburg, 2018 to 2019) in Germany. The 23 inbreds were sowed as replicated checks in an augmented row-column design of other genetic material. The heading time trait was measured as days after planting. Plant height (cm) was measured after heading in Cologne and Mechernich. Seed area (*mm*^2^), seed length (mm), seed width (mm), and thousand grain weight were measured using grains from Cologne, 2017 to 2019, and Quedlinburg, 2018, with a MARVIN seed analyzer (GTA Sensorik, Neubrandenburg, Germany) [50].

### Sequencing

Genomic DNA was extracted using DNeasy Mini Kit Plant (Qiagen). 1 µg of genomic DNA was submitted to mechanical shearing using a Covaris instrument with target fragment length set to 300-500 bp. Library preparation was performed with the NEBNext® Ultra™ II DNA Library Prep Kit for Illumina kit. Methylated adapters were used to prevent conversion of adapters (NEB single methylated index (Catalog # e7535)). Prior to PCR amplification, adapter ligated libraries were bisulfite treated using the EZ DNA-Methylation Lightning™ Kit (Zymo) following manufacturer’s instructions. The resulting paired end libraries were sequenced with Illumina Hiseq2000 and NovaSeq.

### Bisulfite mapping

Raw reads were adapter and quality trimmed with Trimgalore [53] and mapped with Bismark [54] using Bowtie 2 [55], which is recommended especially for large genomes with many repetitive sequences [56].

To increase the mapping efficiency, the reference sequence Morex v3 [57] was SNP corrected with known SNPs for each inbred, respectively. Insertion and deletion correction was also evaluated but the increase in mapping efficiency was less than 0.5 % compared to solely SNP correction (Table S2). This was not enough to justify the added complexity of shifted genomic positions among the genotypes and was therefore not further evaluated.

The Bismark options --score min L,0,-0.6 -X 1200 were selected to optimize the mapping efficiency without sacrificing quality (Table S3). This decreased the minimum alignment score required and increased the maximum insert size of paired end reads [58, 59]. The option -N 1 which sets the number of allowed mismatches to one was also evaluated, but resulted in a worse mapping efficiency.

PCR duplicates were subsequently removed and DNA strands joined using the --comprehensive option.

### Quality control Identification of sample identity

To ensure that none of the samples have been swapped during library preparation, the identity of each sample was verified by comparing SNP calls in the bisulfite data with known SNPs. Firstly, SNPs were identified with Bis-SNP [60] using the uncorrected Morex reference sequence. SNPs were then filtered for shared genomic positions between both datasets. For each shared SNP, the allele calls were compared between the samples. If two samples have the same genotype, they should have the highest number of matching allele calls.

### Conversion rate

Bisulfite conversion rates were assessed using standard protocols utilizing unmethylated chloroplast DNA and counting the number of methylated reads among all reads for each sample [61].

### Reproducibility

In order to assess the technical error of our procedure, Pearson correlations of the proportion of methylated reads between the mixed Sissy inbred line sample and the weighted average across the three separate tissue samples of Sissy were calculated. Only sites with a minimum coverage of 5 in all samples were included.

### DMR identification and characterization

DMRs were called using Methylscore at default settings [62]. Since methylscore does not natively support large chromosomes a customized version was used.

DMRs were classified as TE or genic when they overlapped to at least 50% of their physical length with TEs or annotated genes, respectively. The proportional overlap was calculated relative to the shorter feature so that the shorter length was the dividend. If DMRs intersected both genomic features simultaneously, they were classified as gene + TE. The remaining DMRs were classified as intergenic. The experiment was repeated a second time assessing overlaps of DMRs with specific TE classes present in the barley genome. The number of overlaps was normalized by the respective TE class frequency.

### Population structure analysis

The population structure of the 23 spring barley inbreds was analyzed with Principal Coordinates Analyses (PCoAs) based on euclidean distances of DMRs, SNPs and both expression datasets separately. To quantify the similarities between the population structures derived from the different data sets, a generalized procrustes analysis (GPA) was performed with FactoMineR [63]. Subsequently, 1 − the procrustes similarity indexes were used as dissimilarity measurements in a final PCoA [64]. DMR population structures were calculated separately for each sequence context for the GPA.

### Local association Local association among DMRs and SNPs

To identify DMRs with a significant association to SNPs, the approach of Eichten *et al.* [33] was modified to save computation time. This procedure was applied to all DMRs for the three sequence contexts separately.

Firstly, DMRs were filtered for a maximum of 50% missing data. SNPs were filtered for a maximum of 20% missing data, maximum 20% heterozygosity, minimum minor allele frequency of 5% and to include only biallelic SNPs. Each SNP in the region of any DMR ±10 kb was tested individually, in contrast to the study of Eichten *et al.* [33] with a Wilcoxon rank-sum test, since the residuals were not normally distributed. To define a significance threshold, each DMR that has at least one SNP in the vicinity was tested with 10 regions of 100 random SNPs, which are not in proximity to any DMR. The significance threshold was set to the 5% quantile of all control P values across the three methylation contexts. DMRs were filtered again for at least three significantly associated SNPs. For only those DMRs, 90 additional SNP regions were tested. The significance threshold was revised to the 1% quantile of all control P values. For each DMR that still has at least three significantly associated SNPs, a ranking was conducted between the proportion of associated SNPs in its proximity and the proportion of associated SNPs in each control region. Only when the SNPs in the proximity of the DMR were in the top 5% of the ranking, the DMR was classified as SNP associated.

### Local association among DMRs and gene expression

Spearman’s rank correlation coefficients were calculated between the methylation levels of each DMR and the expression of the closest gene for those DMRs, where the variation of gene expression in inbred lines with the same epiallele was ≥0 for at least one epiallele. Association between DMRs and their closest gene expression state were performed for both expression datasets and the three sequence contexts separately. Benjamini-Hochberg false discovery rate control (FDR) was applied to find significant associations. Additionally, 1000 random DMRs were correlated with 1000 randomly selected genes for each context for comparison.

### Genomic prediction

A linear mixed model was used to analyze each phenotypic trait across all seven environments and estimate adjusted entry means for all barley inbred lines:

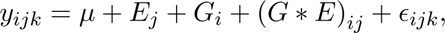

where *y_ijk_* corresponds to the observed phenotypic value of inbred *i* in environment *j* of replicate *k*. The general mean is denoted as *µ*. *G_i_* defines the genetic effect of inbred *i*, *E_j_* the effect of the *j^th^* environment, and (*G* ∗ *E*)*_ij_* the interaction effect of the given inbred and environment. The random error is represented by *ɛ_ijk_*. A variety of predictors was evaluated with respect to their performance to predict the adjusted entry means of all inbreds for each trait measured as the prediction ability (r), a Pearson’s correlation between the observed *y* and the predicted *y*^. The predictors for the genomic best linear unbiased prediction (GBLUP) [65] model were: a Illumina 50K barley SNP array [50, 66], DNA sequencing SNPs [50], seedling gene expression data [47], and DNA methylation data from this study. Both the SNP Array and DNA sequencing SNPs were filtered to remove markers which were monomorphic, had *>*20% missing data, and had a minor allele frequency *<*=5%. Only biallelic SNPs were retained. Missing data were mean imputed. The DNA methylation data was filtered to remove sites with *>*20% missing data, monomorphic cytosine sites and was mean imputed.

*W* for predictor *m* had the dimensions of the number of barley inbreds (*n* = 23) × the number of features of the given predictor (*m_SNP Array_* = 38 285*, m_SNP s_* = 38 725 848*, m_Expression_* = ^43 769*, m*^*Methylation CpG* ^= 238 842 649*, m*^*Methylation CHG* ^= 204 877 046*, m*^*Methylation CHH* ^=^ 789 553 379). All *W* matrices were column centered, standardized to unit variance and denoted as *W* ^∗^. Additive relationship matrices were defined as 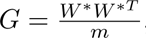, where *W* ^∗^*^T^* was the transposed *W* ^∗^ matrix. To combine the methylation information of the three sequence contexts into one *G* matrix, a joined weighted relationship matrix [67] was formed using the number of cytosines as weights. As the focus of our study lied on the question if methylation can improve genomic prediction using multi-omics predictors, joined weighted relationship matrices of DNA sequencing SNPs and expression; DNA sequencing SNPs and methylation; and DNA sequencing SNPs, expression, and methylation were formed with all possible weight combinations ranging from 0 to 1 in steps of 0.1, where the summation of all weights needed to equal 1 for each combination, respectively.

For the investigation of the prediction ability, 200 five-fold cross-validation runs were used. The median correlation of the five folds was determined and the median of the median correlation across the 200 replicates was calculated as the prediction ability [50].

## RESULTS

The methylation dataset consisted of *>*15.2 billion read pairs which were aligned to a SNP corrected Morex v3 reference sequence with customized parameters to increase the alignment rate (Table S2 and S3). This resulted in *>*10.8 billion unique alignments with a mapping efficiency of 70.8%. The number of uniquely aligned read pairs per sample reached from 152 339 694 to 601 070 430. A negative correlation of -0.39 between the mapping efficiency and the genetic distance of each inbred line and Morex was observed. The average conversion rate across all 26 samples was ≥99%. The coverages of Cytosine sites were on average across all samples 9.4, 9.6, and 10.4 in the CpG, CHG, and CHH sequence context, respectively. Since the main objective of our study was to assess methylation variation as a characteristic of genotypes instead of different tissues, Pearson correlation coefficients between a pool of three Sissy tissues and the average of the three individual tissues was calculated. The correlation coefficients were 0.86, 0.89, and 0.66 for the CpG, CHG, and CHH sequence context, respectively (Figure S1). This approach also served to assess the reproducibility of the methylation levels in this study.

### The barley methylome

The average methylation levels across the 23 barley inbred lines were 88.6%, 58.1%, and 1.4% in the CpG, CHG, and CHH sequence context, respectively. Mean methylation levels calculated in bins of 5 Mbp across 23 inbred lines were tending to be lower at the distal regions of the chromosomes in the CpG and CHG context where the repeat content was also lower (Figure 1). For CHH methylation, the opposite trend was observed. CpG methylation levels showed local minima in the centromeric region, while CHH methylation showed a minor peak in that region. The overall shape of the curves was largely uniform across the chromosomes with a mean standard deviation of 8.71%, 10.71%, and 0.28% in the CpG, CHG, and CHH context, respectively (Figure S2).

**Fig. 1:**
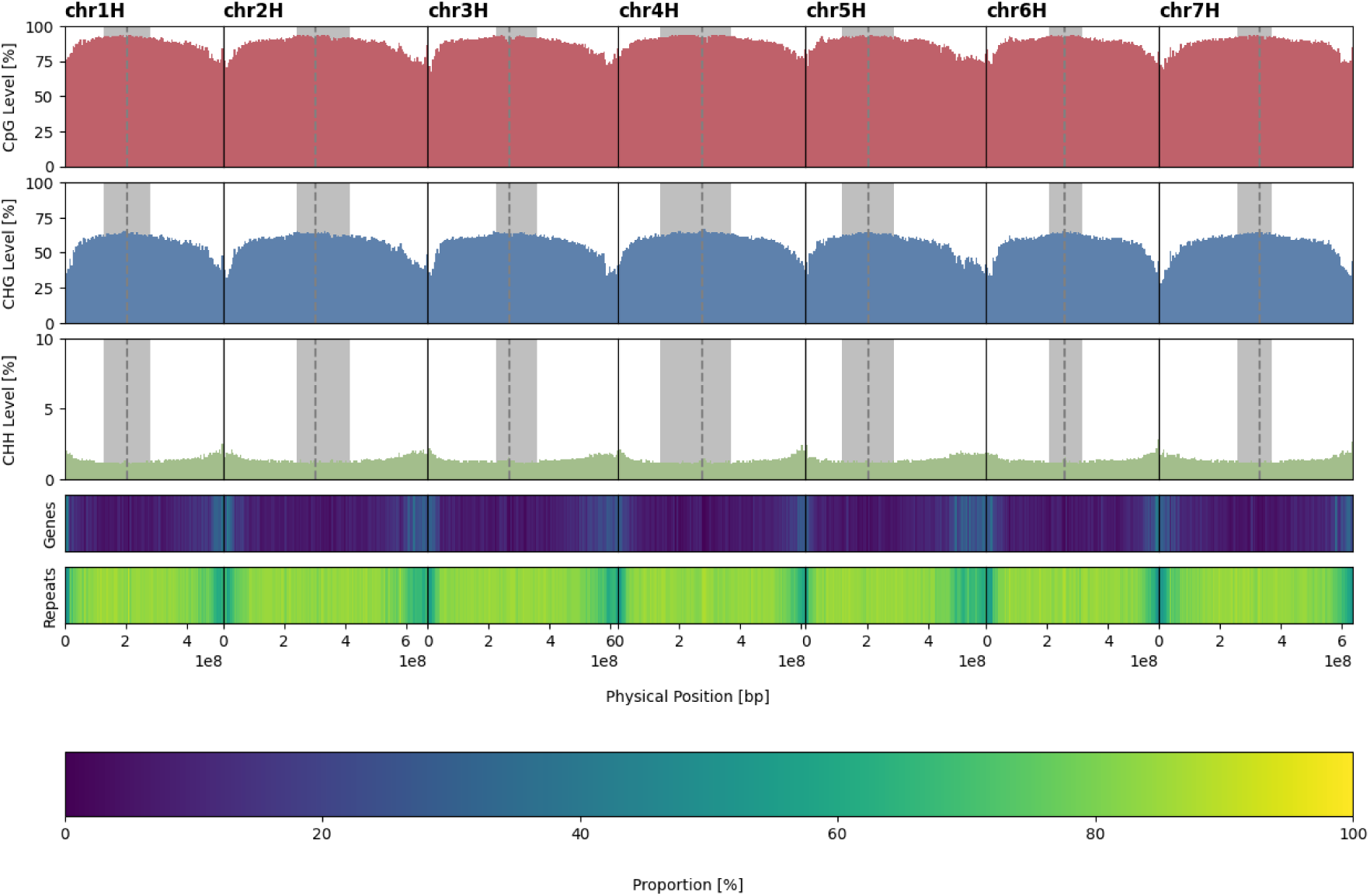
Mean methylation levels of the three sequence contexts calculated in bins of 5 Mbp across 23 inbred lines. The dashed gray lines indicate the centromere with the pericentromeric regions highlighted in gray [48]. Proportions of genes and repeats were extracted from the Morex v3 GFF [57].

In order to assess DNA methylation on a local level around genes, these were averaged in 100 bp bins between 10 kb up- and downstream of all TSS and TTS and then averaged across the inbreds (Figure 2). In all contexts, DNA methylation minima were observed at TSS and TTS. The slope of methylation was generally more steep on the side before a TSS and after a TTS compared to after a TSS and before a TTS. The trend of the curves at TSS and TTS was largely symmetric. CpG and CHG methylation levels varied strongly between the genotypes around TSS and TTS, but the overall shape of the curves showed the same trend. CpG methylation displayed, in comparison to the other two sequence contexts, the steepest decline and increase right before and after a TSS and the steepest increase after a TTS. CHG methylation showed a less steep increase over a long genomic sequence after a TSS in comparison to CpG methylation. Interestingly, CHH methylation appeared to have a peak right before a TSS, followed by a minimum before returning to the normal level. The opposite trend was observed at TTS. However, the peak in CHH methylation was more pronounced before a TSS than after a TTS.

**Fig. 2:**
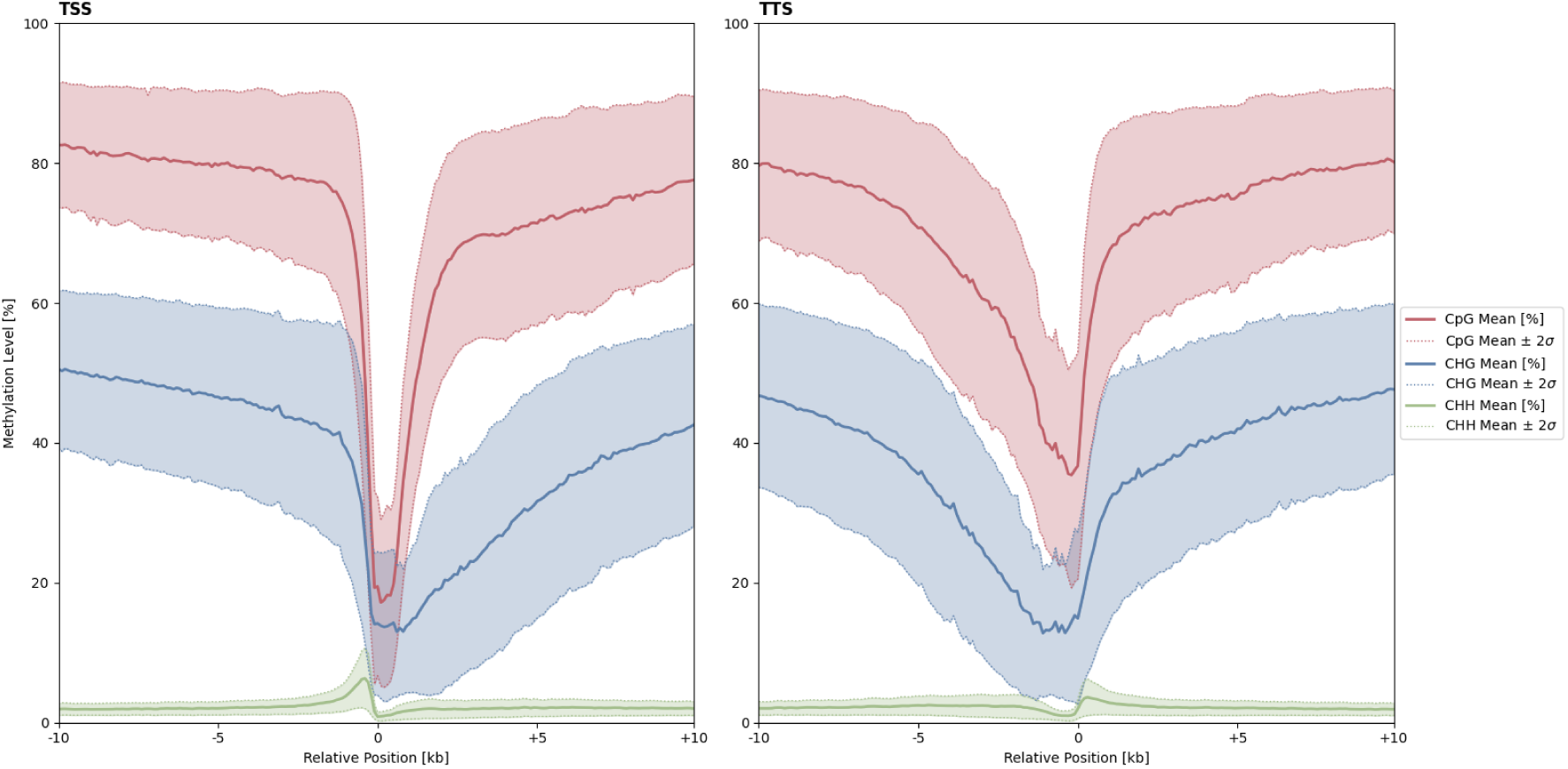
Mean methylation levels of 10 Kbp upand downstream regions of transcription start sites, left and transcription termination sites, right calculated in bins of 100 bp across 23 inbred lines. Variability among the inbreds is illustrated by an approximation of the 95% confidence interval for each sequence context as the colored area.

### Gene methylation and expression

Genes of the leaf expression dataset of Sissy were categorized based on their expression level in five groups (100% quantile ≥ High *>* 75% quantile; 75% quantile ≥ Medium High *>* 50% quantile; 50% quantile ≥ Medium Low *>* 25% quantile; 25% quantile ≥ Low *>* 0% quantile; None = 0) and divided together with their corresponding 2 kb up- and downstream regions into 200 bins, respectively, across their physical length to assess methylation levels among genes of different expression levels in one genotype. Subsequently, average methylation levels were calculated for each bin in the Sissy leaf tissue bisulfite data (Figure 3). Highly expressed genes were strongly methylated in the CpG context and moderately methylated in the CHG context. They were also only slightly more methylated in the CHH context compared to less expressed or silenced genes. On the other hand, CpG and CHG methylation in the up- or downstream regions of the gene body were associated with a low expression. This was in contrast to CHH methylation, which was positively associated with an increase of expression. Interestingly, silenced genes showed the highest CHG methylation levels in the gene body region, while highly expressed genes were only slightly less methylated. The same results were obtained using the Sissy seedling gene expression and methylation data (Figure S3).

**Fig. 3:**
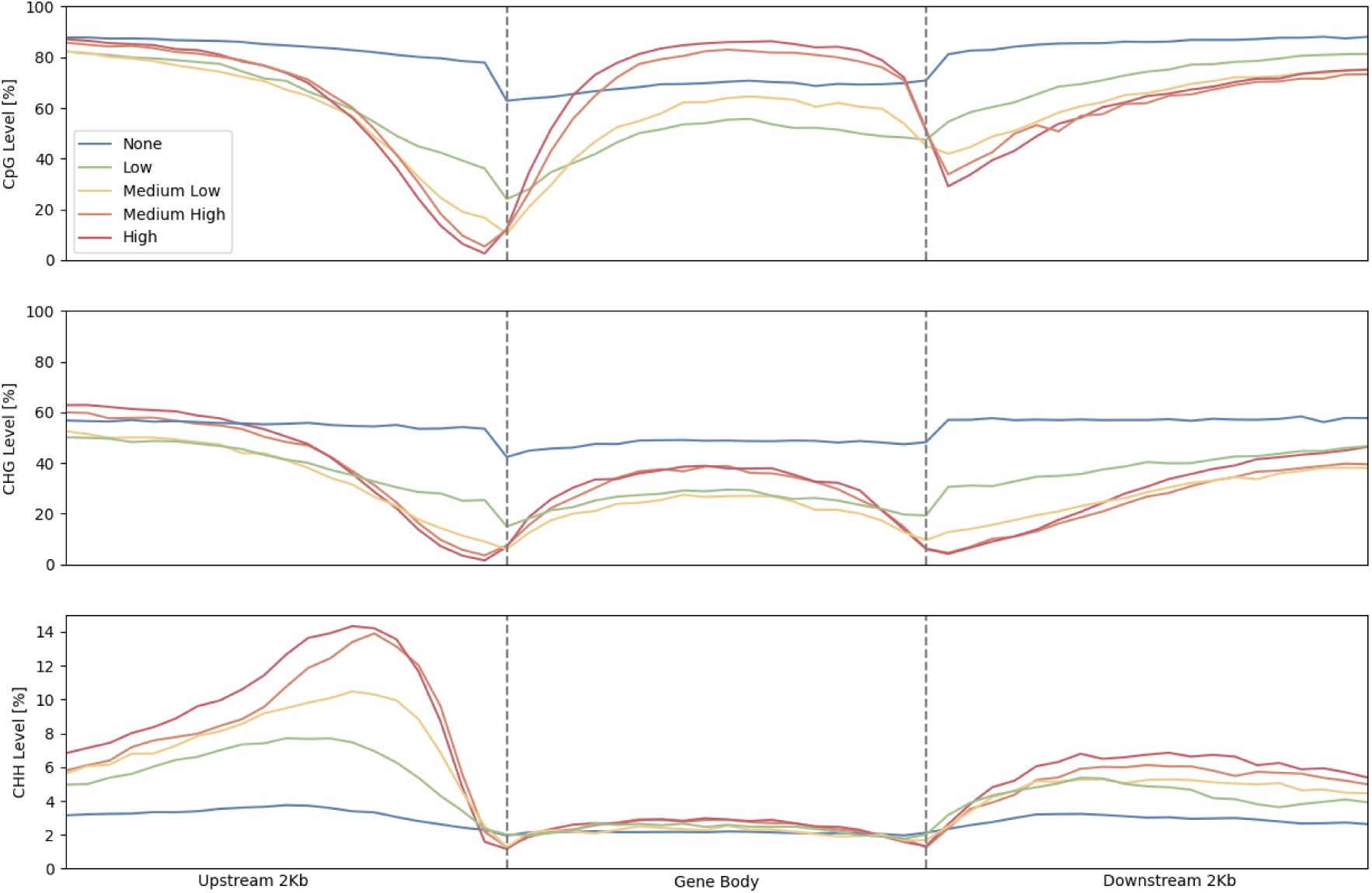
Average methylation levels of genes in Sissy’s leaf tissue categorized based on their expression levels in the same tissue (100% quantile ≥ High *>* 75% quantile; 75% quantile ≥ Medium High *>* 50% quantile; 50% quantile ≥ Medium Low *>* 25% quantile; 25% quantile ≥ Low *>* 0% quantile; None = 0).

### Differential methylation

Among the 23 barley inbred lines, 244 689, 151 992, and 103 115 DMRs with a minimum of five Cs per DMR in the CpG, CHG, and CHH context were discovered, respectively. Even though CHH sites were about 2.1 times more common than CpG and CHG sites together, they showed the least amount of differential methylation with a proportion of 20.63% of all DMRs. DMRs tended to be located at the ends of the chromosomes with up to 22.78 fold more DMRs compared to the pericentromeric regions (Figure 4a). This was especially noticeable towards the 3’ end. Nevertheless, a local maximum in the number of DMRs was found right at the centromere in all chromosomes when considering all three sequence contexts. DMRs of the CHG and CHH context were distributed similarly to that of the CpG contexts. However, CHH DMRs not only tended to be fewer, but were also generally shorter than DMRs of the other two contexts (Figure 4c).

**Fig. 4:**
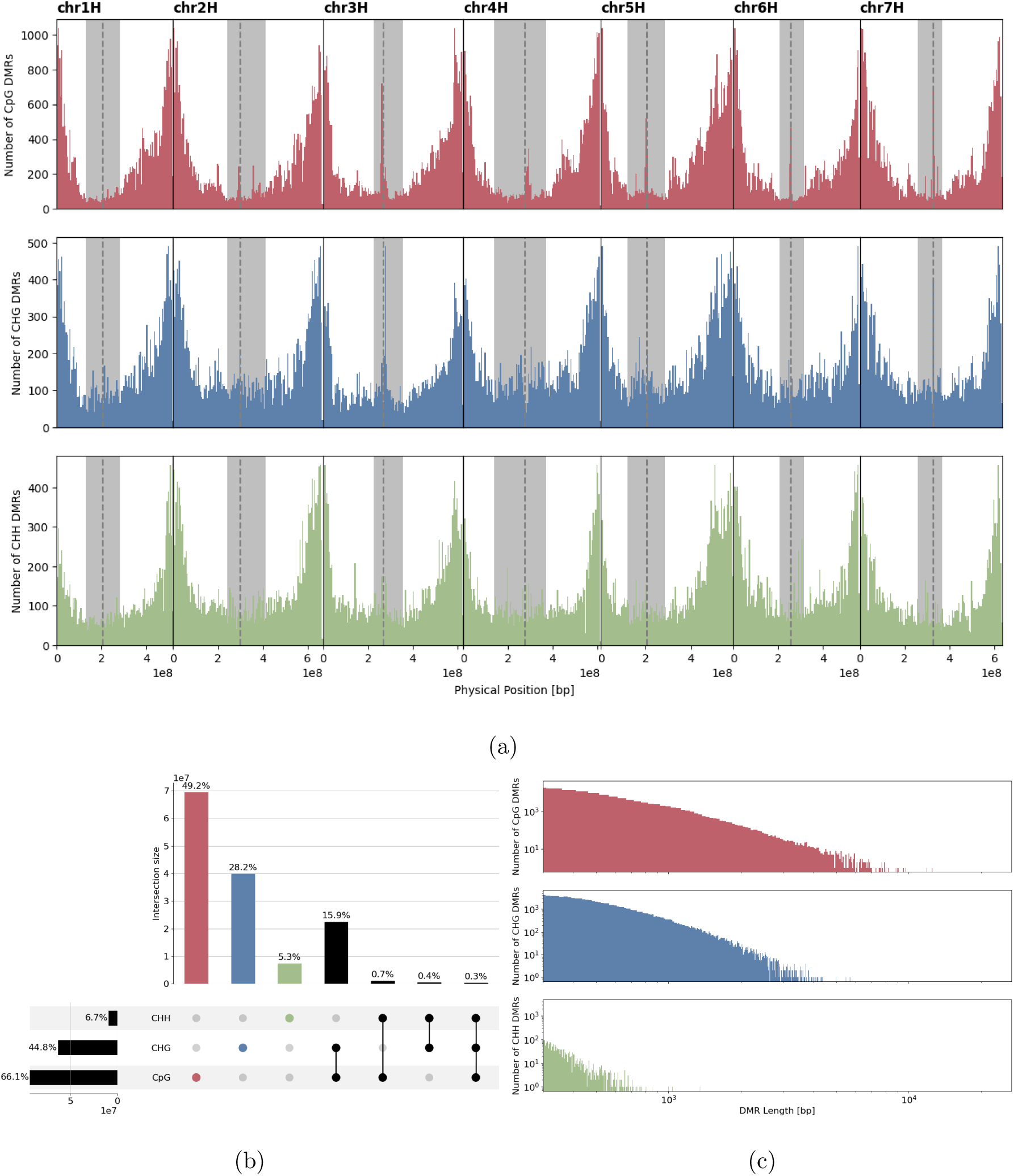
a) Distribution of differentially methylated regions (DMRs) in bins of 5 Mbp across the chromosomes. The dashed gray lines indicate the centromeres with the pericentromeric regions highlighted in gray. b) Proportional overlap of physical positions between DMRs of all three methylation contexts. c) DMR length distribution on a logarithmic scale.

In addition to the correlation of methylation levels between the physically bulked Sissy tissues and the mean of the separate tissues, also DMRs between the three tissue samples of Sissy were called. In comparison to the mean of 100 replications of three randomly selected unique combinations of barley inbred lines, the amount of DMRs between the three tissues were about 224, 24193, and 2 times lower in the CpG, CHG, and CHH context, respectively (Table 1).

**Table 1:**
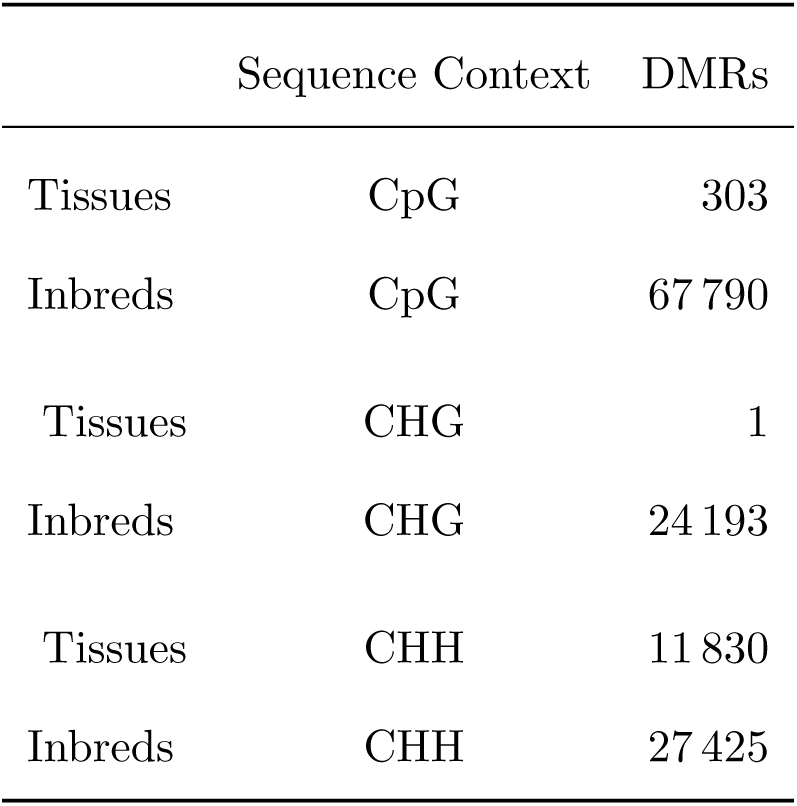
Summary of the number of differentially methylated regions (DMRs) between three tissues of inbred Sissy and the average of 100 random unique combinations of three inbred lines.

DMRs were classified as TE, genic, gene + TE, or intergenic based on positional intersections (Figure 5a). The majority of all DMRs were classified as TE in the CpG and CHH context with a share of 45% and 70%, respectively. Intergenic was the second most common category with 40% and 25% for the CpG and CHH context, respectively. In contrast, CHG DMRs were most often classified as intergenic with a share of 48% and TE was the second most common class with 40%. For CpG, the highest proportion of genic DMRs across the three sequence contexts was observed with 13%. Repeating the experiment with no minimum required overlap-size did not change the results considerably. Since the majority of DMRs were classified as TE, a more detailed analysis was conducted to determine which TE classes most frequently overlapped with DMRs. In total as well as in the CpG context, intersections with retrotransposons of the RXX class were the most common (Figure 5b). In the CHG sequence context, intersections with transposons of the DHH class were more frequent, while DTX transposon intersections were the most common in the CHH context. Additionally, DMRs were found to be overrepresented in miniature inverted-repeat transposable elements (MITEs) assessed with a binomial test (P *<* 0.01) in all three contexts.

**Fig. 5:**
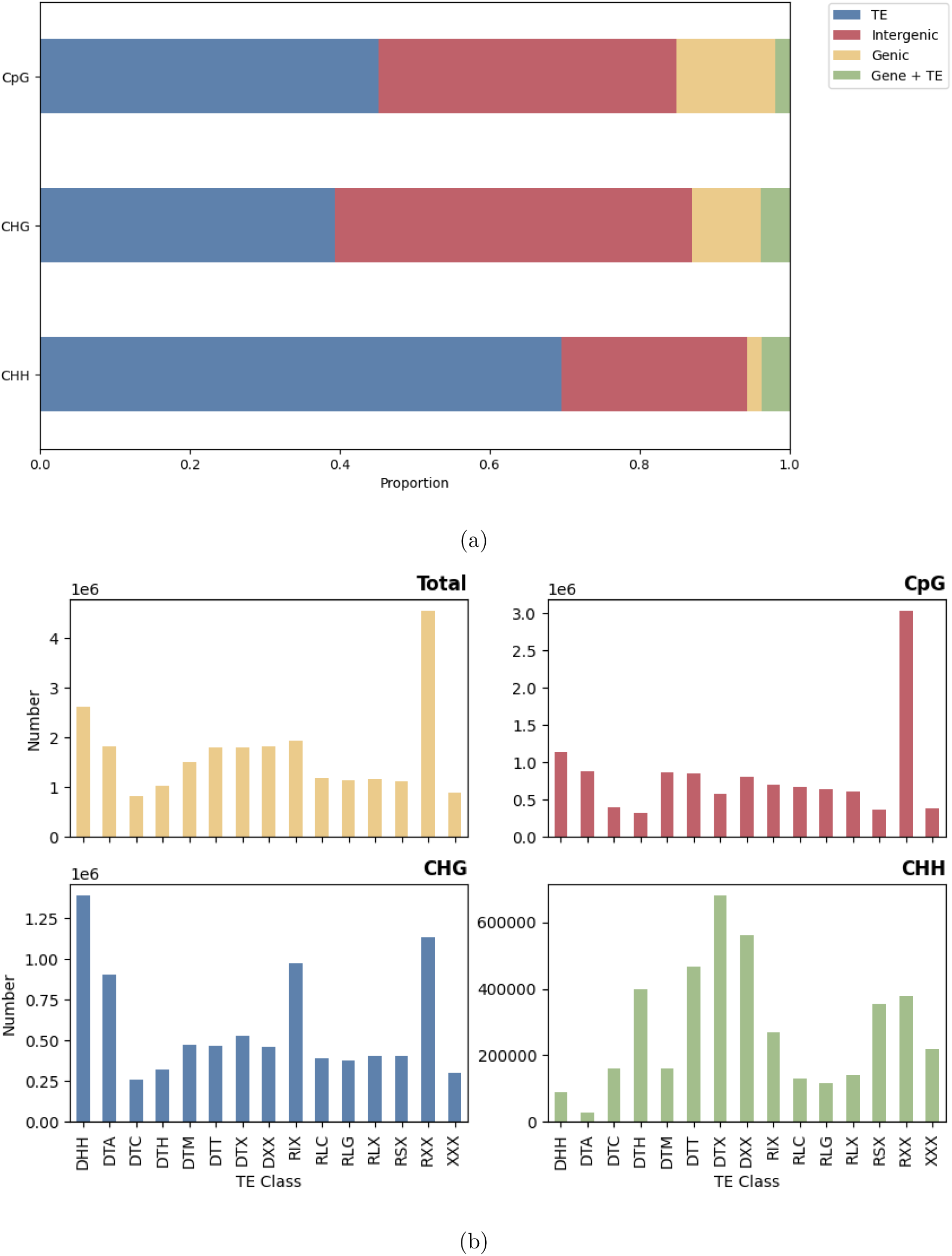
a) Annotation of differentially methylated regions (DMRs) intersecting with genic, transposable element (TE), intergenic, or both TE and genic features for each sequence context. b) Number of DMRs overlapping with specific TE-classes normalized by the frequency of the given TE-class in the barley genome.

### Population structure analysis

Euclidean distances were calculated from DMRs and subjected to a PCoA (Figure 6a). The axes explained 11.48% and 7.54% of the total variance. The resulting population structure formed three clusters. One cluster consisted of two-row landraces and cultivars, while six-row inbred lines were clustered in a separate cluster. The cluster in the center of the PCoA consisted out of two- and six-row inbred lines. Inbred lines from the same country of origin mostly fell into the same cluster like the Syrian IG31424 and HOR12830, the Turkish HOR7985 and HOR8160, or the Indian Lakhan and Kharsila.

**Fig. 6:**
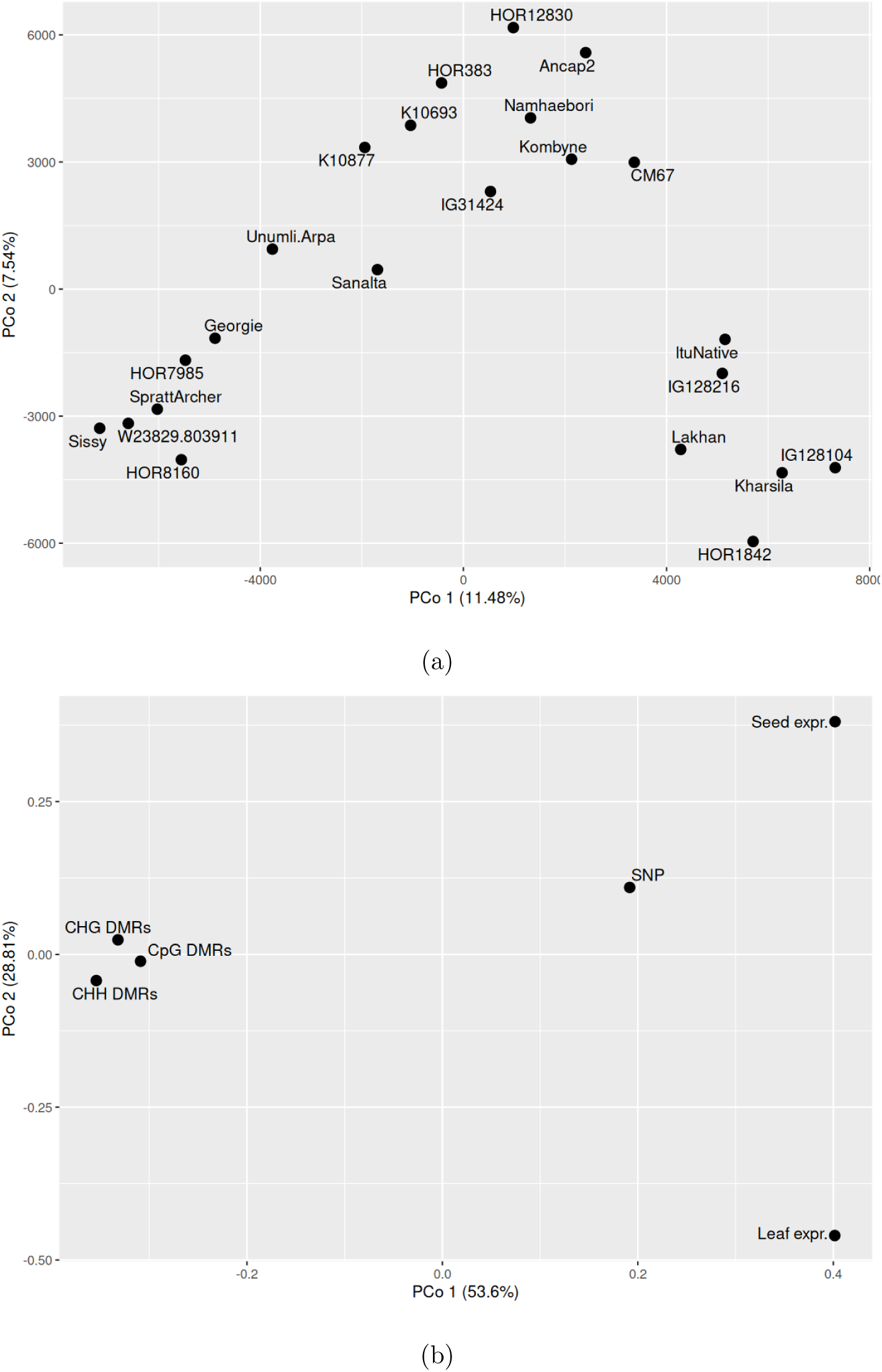
a) Principal coordinate analysis (PCoA) of 20 inbred lines based on euclidean distances determined by differentially methylated regions (DMRs). The percentage values on the axes refer to the proportion of explained variance by the respective axis. b) PCoA comparing the information content of CpG DMRs, CHG DMRs, CHH DMRs, single nucleotide polymorphisms (SNPs), and expression based on a generalized procrustes analysis.

Comparing the population structures derived from SNPs, the two gene expression datasets and DMRs, separate for each context, using GPA, revealed considerable differences (Figure 6b). The DMR based population structures clustered closely together, while the SNP and expression population structures were spread out. Interestingly, the distances between the SNP and expression population structures were smaller between each other than to the DMRs. Furthermore, we observed a correlation of -0.07 between the distance matrices calculated from SNPs and DMRs using a Mantel test with 99 permutations.

### Local association of DMR

In the CpG, CHG, and CHH sequence context, 36.37%, 36.20%, and 16.59% of the DMRs were significantly (*p <* 0.01) associated with at least three local SNPs, respectively. SNP associated DMRs largely accumulated at the ends of the chromosomes in all three sequence contexts, notably towards the 3’ end (Figure S4). This trend was similar to the distribution of all DMRs with an average Pearson’s correlation coefficient of 0.91 across all chromosomes and sequence contexts (Figures 7 and 4a). In addition, we observed that the Kolmogorov-Smirnov test of uniformity of the number of SNP associated DMRs corrected for the number of DMRs per bin was not significant (*p <* 0.05) for all chromosomes and contexts. This shows, that when corrected for the total amount of DMRs, the SNP associated DMRs distributed more towards the far chromosome ends than the total amount of DMRs. Most notably, some chromosomes showed local maxima in the proportion of SNP associated DMRs among all DMRs in the pericentromeric region, while others show local minima even when corrected for the total amount of DMRs (Figure 7).

**Fig. 7:**
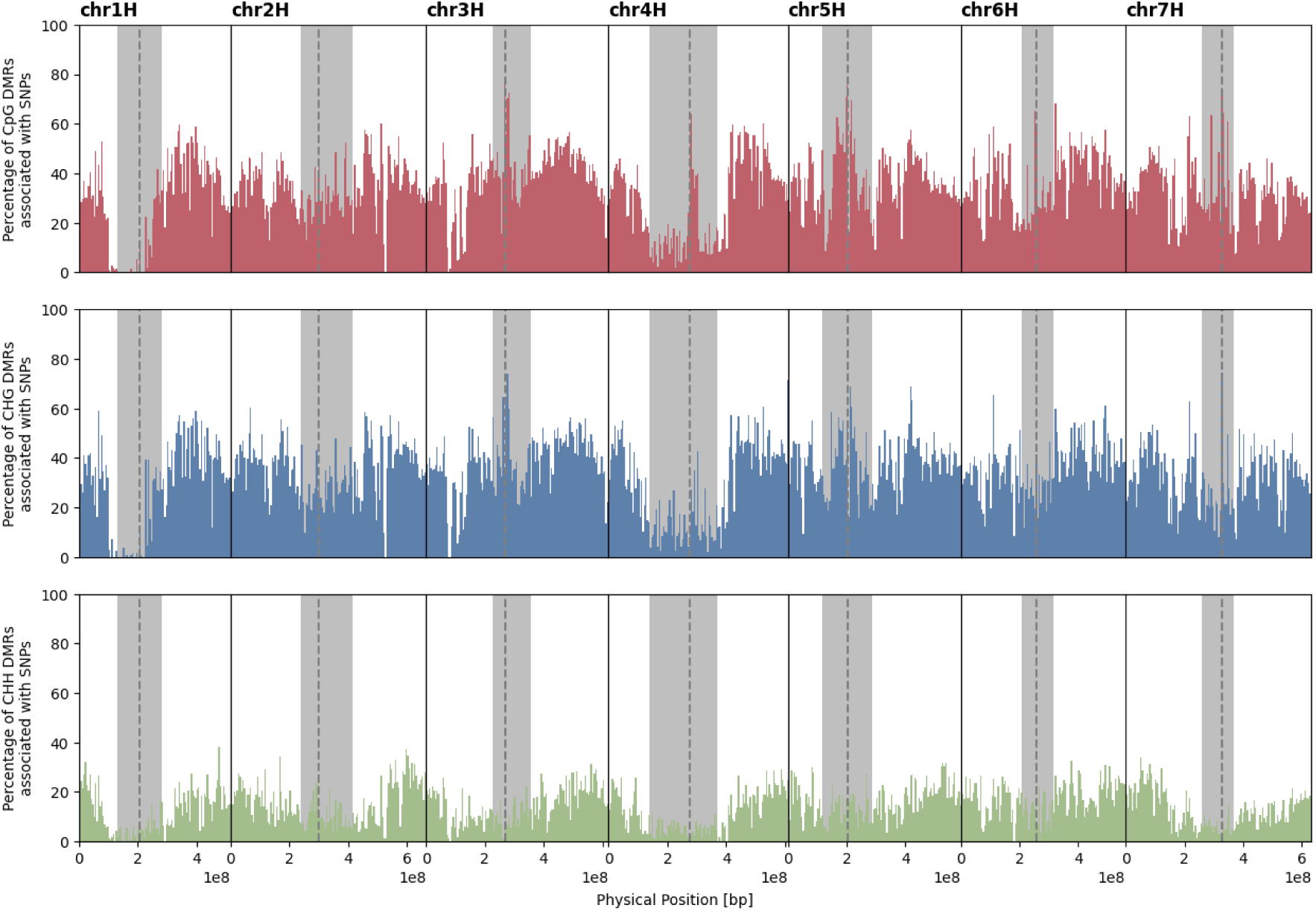
Percentage of single nucleotide polymorphism (SNP) associated differentially methylated regions (DMRs) among all DMRs in bins of 5 Mbp across the genome. The dashed gray lines show the middle of the centromere with the pericentromeric regions highlighted in gray.

The distribution of the Spearman’s correlation coefficients between the methylation level at DMRs and the gene expression of the closest gene illustrated that only a low proportion of all genes was highly positively or also negatively correlated with methylation across the inbreds, while most genes showed low to medium correlations in all sequence contexts (Figure S5). The same was also the case for 1000 random DMR correlations with 1000 randomly selected genes. Except for CpG and CHG, where a negative trend of correlation between gene expression and methylation at DMRs was found in close proximity to the TSS, all distributions were symmetric around 0 across the various distance groupings and did not differ from the random correlations. Only if significant correlations (FDR *<* 0.05) were considered, the distributions showed a negative correlation trend in the CpG and CHG contexts upstream of, and most notably at, the TSS (Figure 8). Correlations downstream of the TSS tended to be positive in the CpG context, while they were strongly negative in the CHG context up to 2kb from the TSS. In the CHH context, all correlations between the DMRs and gene expression were strongly positive regardless of the distance to the TSS. However, a slight pattern can be recognized in stronger positive correlations closer to the TSS. Spearman’s correlation coefficient distributions of all contexts were similar between the leaf and seedling expression datasets.

**Fig. 8:**
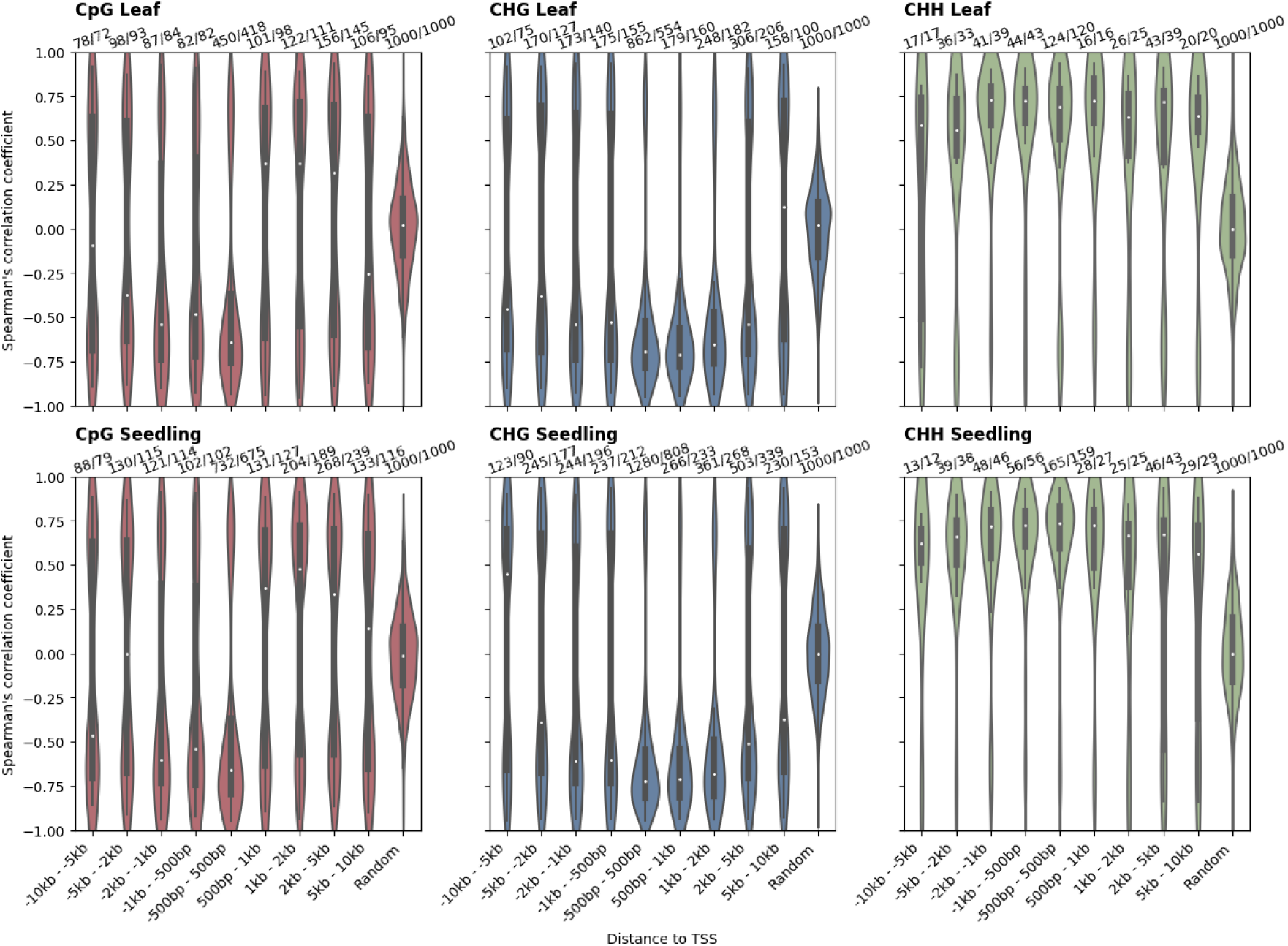
Spearman’s rank correlation coefficient distributions of differentially methylated regions (DMRs) with gene expression filtered using Benjamini-Hochberg’s false discovery rate control (FDR*<*0.05) and grouped by the DMR’s distance to the respective transcription start site (TSS) ±10 kb in intervals of 5 kb to 500 bp for the leaf and seedling expression data separately. Additionally, 1000 DMRs were randomly correlated with 1000 random genes, presented in the last category of each subplot. The numbers above the categories reflect the number of correlations, as well as the number of unique genes in the respective category.

### Association of DMR and phenotypic variation

One of the CHG DMRs with a high correlation of its methylation level with gene expression of +0.53 with one gene in the seedling expression dataset was indentified in an intronic region of *VERNALIZATION1* (*VRN-H1* ; Figure 9). *VRN-H1* transcribes a MADS-box transcription factor which promotes flowering and is part of the main genes regulating vernalization response [68]. We observed two epialleles at this DMR: epiallele 1 with a mean CHG methylation level of 73% and epiallele 2 with a mean CHG methylation level of 52% (Figures 9a and 9b). The inbred lines carrying epiallele 1 showed an increased expression of *VRN-H1* in both expression dataset and an earlier flowering time compared to the inbreds carrying epiallele 2 (Figure 9c). The DMR was not associated with SNPs or structural variants in the regulatory or coding region of *VRN-H1* [50]. Methylscore was unable to assign an epiallele to inbreds HOR1842, IG128104, and Sissy, which showed an average CHG site coverage of 1.3, 2.6, and 1.6 within the DMR, respectively.

**Fig. 9:**
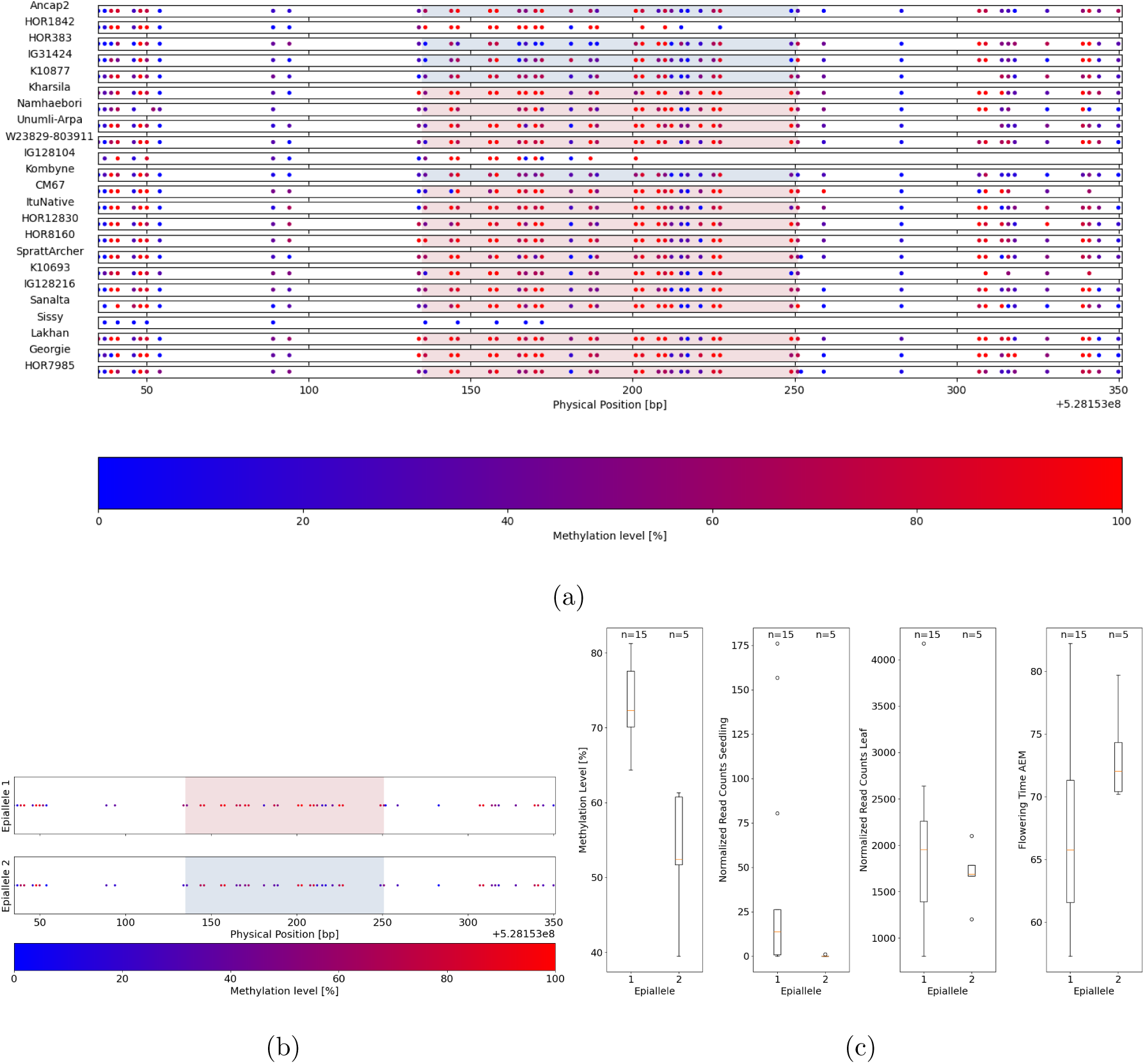
a) Differentially methylated region (DMR) in an intronic region of *VRN-H1*. The DMR is highlighted with a colored background, where the inbred lines carrying the same epiallele share the same color. The red background corresponds to epiallele 1 with a mean methylation level of 73% and the blue background corresponds to epiallele 2 with a mean methylation level of 52%. For inbred lines with a white background color no epiallele could be assigned by Methylscore. The dots represent CHG sites colored according to their methylation level. The inbred lines are sorted by their expression of *VRN-H1* in ascending order. b) Average methylation level per CHG site for both epialleles. c) Box plots of the CHG methylation level at the DMR, gene expression of *VRN-H1* in the seedling dataset, gene expression of *VRN-H1* in the leaf dataset, and adjusted entry means (AEM) for flowering time for inbred lines carrying either epiallele 1 or 2, respectively.

### Methylation as a predictor of phenotypic variation

A SNP array, DNA sequencing SNPs, gene expression, and methylation data were used as single predictors to assess the median prediction ability for six traits in a GLUP framework. In addition, for the DNA sequencing SNPs, gene expression and methylation data, joined weighted relationship matrices were created with all possible weight combinations between 0 and 1 in steps of 0.1 to select the one with the highest median prediction ability (Figure 10). The prediction abilities across all six traits ranged from 0.17 to 0.85. The joined relationship matrices of DNA sequencing SNPs, gene expression, and methylation (S+M+E) and the ones of DNA sequencing SNPs with methylation (S+M) had the highest prediction ability across the six traits followed by the SNP array and the DNA sequencing SNPs alone. The mean optimal weight across all traits in the S+M scenario was the highest for methylation with 0.63. In the S+E scenario, DNA sequencing SNPs had the highest mean optimal weight of 0.83. The highest mean optimal weight in the S+E+M scenario was observed for methylation with 0.52.

**Fig. 10:**
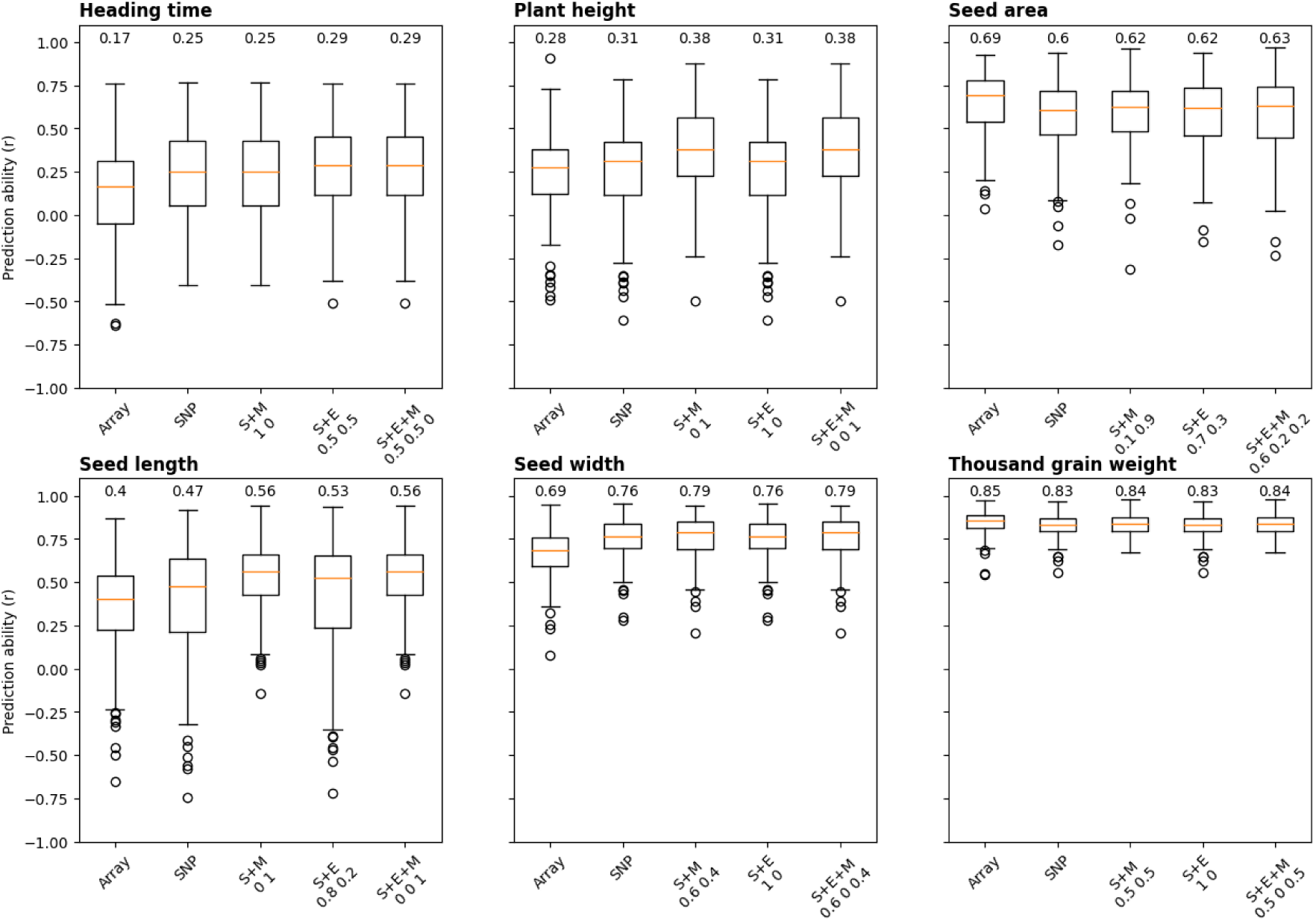
Prediction abilities for 6 traits from 200 five-fold cross-validation runs using either a single nucleotide polymorphism (SNP) Array, DNA sequencing SNP, combined SNP and methylation (S+M), combined SNP and gene expression (S+E), or combined SNP, methylation and gene expression (S+E+M) to establish relationship matrices among 23 barley inbred lines. The numbers below the combined datasets show the weights for the joined weighted relationship matrix with the highest prediction ability.

For the traits plant height and seed length, methylation outperformed any other predictor, which was indicated by a weight of 1 in the optimal joined relationship matrices. The prediction ability of plant height and seed length was improved by 0.1 and 0.16, respectively, using methylation data as a predictor compared to the SNP array.

For the traits seed area, seed width, and thousand grain weight, the combination of S+M outperformed the combination of expression data and DNA sequencing SNPs (S+E) and DNA sequencing SNPs alone with improvements ranging from 0.01 to 0.03 compared to the DNA sequencing SNPs. Instead, the prediction ability for the traits seed area and thousand grain weight was the highest using the SNP array. The prediction ability could not be improved by adding methylation data for the prediction of the trait heading time in comparison to S+E.

## DISCUSSION

From the observed high conversion rates of *>*99%, which are comparable to those reported in literature [69, 70], it can be concluded that the methylation data for all inbred lines is of high quality and that there is no bias due to the bisulfite treatment. In addition, we observed high and significant correlation coefficients among the technical replicates which indicates a good reproducibility of this experiment in the CpG and CHG context. In contrast, for the CHH context, the reproducibility was only medium (Figure S1). Furthermore, this observation of a high correlation between the methylation rates of a bulk sample and an artificial bulk sample created from individual tissue sequence data suggested that the bulking of tissue samples is a reasonable approach to assess a genotype specific methylation profile at moderate costs. This conclusion is also supported by the observation of several orders of magnitude less differential methylation among the tissues, especially in the CpG and CHG context, compared to the amount of differential methylation among the inbreds (Table 1). This illustrates that it is possible to investigate methylation differences of inbred lines even if the samples consisted of different or mixed tissues.

In our study, no full genome sequence was available for all 23 inbreds. Instead, we have used the SNPs of whole genome sequencing of the 23 inbreds to create a SNP corrected reference sequence from the typically used barley reference sequence Morex [57]. This approach, however, did not fully remove the mapping bias. We observed a negative correlation between the mapping efficiency and the genetic distance of each inbred line and Morex of -0.39. However, as only sufficiently covered cytosine sites are considered for differential methylation analyses and the lowered mapping efficiency of distant inbreds effects both methylated reads and unmethylated reads equally, we expect that no methylation level bias is introduced by the differing relatedness of the characterized inbreds to Morex.

### The barley methylome

The average methylation level of barley was exceptionally high when comparing it to *A. thaliana* or rice [71, 72]. Compared to maize, the average methylation levels of barley in the CpG and CHG context were about 10% higher and 1% lower in the CHH context [73]. The higher methylation levels in the CpG and CHG context can be explained by barley’s large genome with many repetitive elements (Figure 1). However, these inter species comparisons should be treated with caution as the library preparation as well as the analysis methods in our study are different from the studies mentioned above. Furthermore, the results for the CpG and CHG context are in line with a previously introduced RRBS study in barley [44]. However, the average CHH methylation level observed in our study was about 48% lower compared to the study using the RRBS approach. This might be due to sampling bias as the previously mentioned study employing RRBS just covered 0.7% of the barley genome, whereas we used the gold standard approach of WGBS (cf. Yong *et al.* [74]).

On a genome level, CpG and CHG methylation levels largely followed the distribution of the repeat content with correlation coefficients of 0.67 and 0.82 (data not shown, Figure 1). Interestingly, barley showed a slight enrichment of CHH methylation around the centromere like *A. thaliana* and contrary to rice [75]. Previous research in *A. thaliana* has shown that centromere methylation plays an essential role in chromosomal stability and regulates the frequency of meiotic recombinations [76–78].

On the level of genes, the steep drop in methylation level in the CpG and CHG context at TSS and TTS is in agreement with previous studies [44, 79]. The peaks in CHH methylation before a TSS and after a TTS have originally been described in maize as CHH islands [80] and were previously assessed in barley as well as other grass species [81], where they were associated with genes with a low gene body methylation. In the study of Martin *et al.* [81], genes with a low gene body methylation and CHH islands had a generally higher expression than genes with only one of these features.

The approximation of the 95% confidence interval of methylation across inbreds indicates that the course of the methylation level around genes is the same among the inbred lines. However, this analysis also suggested the presence of large differences in the extend of methylation among inbred lines (Figure 2), which will be discussed later.

### Gene methylation and expression

On the level of a single inbred line, a clear relationship between the methylation levels of genes and their expression was observed (Figure 3 and S3). Our observation indicated a high potential to predict the expression of genes based on methylation level changes when considering one genotype. Promoter methylation in the CpG and CHG context represses gene expression, while CHH methylation in the upstream 2kb region increases gene expression on the genotype level, which is in line with previous research in barley and other crops [34, 43]. The highly expressed genes had the highest methylation level in the gene body in the CpG and CHH context, however in the CHG context, the most methylated genes were not expressed, which is also observed in the previous study [43]. These observations could potentially be explained by a high correlation of CHG methylation and dimethylation of histone H3 lysine 9 (H3K9me2), a histone modification that typically leads to transcription silencing. This aspect, however, is debated [82, 83] and requires further investigation. These findings highlight the importance of accounting for the methylation context as well as the location of the DNA methylation when investigating its influence on gene expression.

### Differential methylation

Explaining the variation of plant phenotypes is one of the key aspects in plant genetics [84]. However, relying on sequence variation alone has resulted in missing heritability in previous research [3]. As suggested by the tight relationship between methylation and gene expression within one genotype, investigating the differences in DNA methylation may be one of the key aspects to unravel more of the missing heritability [3, 4].

More CpG DMRs were identified than CHG DMRs (Figure 4a). The CHH context shows the least amount of DMRs despite being the most frequent site (Figure 4a). This can probably be accounted to CHH sites being largely unmethylated (Figure 1). The accumulation of DMRs towards the chromosome arms that was observed in our study (Figure 1) might be explained by methylation being less conserved in the euchromatic chromosome arms between the inbred lines e.g. epimutation rates are more prevalent in euchromatic regions with many genic and intergenic sequences and depleted in pericentromeric regions [85]. A potential explanation might be that errors during meiosis lead not only to structural variants [86] but also differences in methylation. This hypothesis is supported by a very high average correlation between the amount of DMRs and the recombination rate of 0.76 across all chromosomes and contexts (data not shown, Casale *et al.* [87]). However, why this finding can not be observed in soybean and is either not present in maize or only to a far lower extent requires further investigation (Xu *et al.* [34], Shen *et al.* [88], and Li *et al.* [89]; Figure 4a).

Additionally, we characterized the DMRs based on their overlaps with genic, intergenic or TE features. CpG DMRs were located more frequently at annotated genes than DMRs of the other contexts. This is likely because genes are predominantly CpG methylated [21].

DMRs of all three sequence contexts largely targeted TE, especially of the RXX class, which is unsurprising given their high methylation ratio and the overall high repeat content in the barley genome (Figure 5a and 5b). This association of DMRs and TEs was also observed in other crops, as DNA methylation is a key factor regulating their expression [88–90]. Additionally, the over-representation of DMRs at MITEs, a known TE superfamily involved in post transcriptional regulation of gene expression [91–93], of all methylation contexts may hint that DMRs serve as an additional layer of gene expression regulation that activates or deactivates MITE transcription.

We observed local maxima of DMRs at the centromeres that might be explained by sequence variation among the studied inbred lines. In rice, functional centromeres were either hyper- or hypomethylated compared to the pericentromeric regions, which may be the result of sequence variation as hypothesized by Yan *et al.* [94]. This means, depending on the sequence composition of the inbred line, different centromeres can either have a higher or lower methylation level compared to the pericentromeric regions in the same cultivar. As local maxima of sequence variation between the same barley inbred lines were previously reported at centromeres [50], it may be possible that in our study the sequence variation among the inbred lines is a cause for the local maximal of DMRs at the same centromeres. This hypothesis is supported by the high amount of differential methylation that targeted repeat rich genome regions (Figure 5a) but still requires further research.

### Linkage disequilibrium of DMRs and SNPs

Investigating the linkage disequilibrium of DMRs and SNPs is crucial to understand the independence of epigenetic variation to sequence variation as only such variation is able to explain the missing heritability. In barley, 32% of all DMRs across all three contexts were associated with SNPs (Figure 7 and S4). Similar results were observed in other crops like maize with less than 40% SNP associated DMRs and soybean with about 23% [34, 88]. We observed that SNP associated DMRs distributed more towards the far chromosome arms than the total amount of all DMRs (Figure 7). However, unraveling the reasons of the local minima of SNP associated DMRs in the pericentromeric regions of some chromosomes, while others show local maxima, requires further research.

The low amount of SNP associated DMRs highlights that there is a considerable difference in information content derived from DNA methylation compared to sequence variation. This hypothesis is also supported by the GPA of population structures derived from DMRs and SNPs (Figure 6b), where these differences in information content can be observed. This is further supported by the observation of a correlation of -0.07 between the distance matrices calculated from SNPs and DMRs. A low level of linkage disequilibrium between SNPs and DMRs emphasizes that DMRs reveal new levels of genotypic information not accessible by other means. As we observed a tight relationship between the gene expression and the extent of methylation within one genotype, as previously discussed, we were interested in understanding the predictive power of methylation differences among inbreds on the respective gene expression.

### Association of DMRs with gene expression

The 23 inbred lines showed large differences in their methylation level around TSS and TTS (Figure 2). We observed that most DMRs at genic loci were of the CpG context (Figure 5a), as described previously. However, when looking at the number of significant associations of DMRs with the expression of the closest gene, most of them were in the CHG context (Figure 8). This highlights that the relevance of CHG methylation for the regulation of gene expression, at least in barley, is higher than previously described in literature [83, 95]. This can also be observed for the gene *VRN-H1*, which regulates the vernalization response in barley (Figure 9).

When considering the association of DMRs with the expression levels of adjacent genes, it can be concluded that these differences are having only minor effects on gene expression (Figure S5). Only when considering significant associations, strong correlations between DMRs and gene expression were observed (Figure 8). However, significant associations were only observed for 5.69% out of all annotated genes across the 7 chromosomes, which is of the same order of magnitude as the 3.38% observed gene-DMR associations in maize [34]. The percentages of DMRs with gene expression associations were 1.3% and 1.7% for the leaf and seedling expression dataset, respectively. This illustrates, that the effect of methylation on gene expression across inbred lines is limited and suggests that other cis-[96] or possibly trans-effects [97] are responsible for the differential expression of the majority of genes. However, this requires further research.

As described above for barley (Figure 3) but also earlier in literature [21, 43], a pattern of promoter methylation in the CpG and CHG context leading to downregulation of expression emerges when considering different genes in a singular genotype, while CHH methylation results in the contrary. When observing the methylation differences and gene expression variation of the same genes across a diverse set of inbreds instead, a similar, however more specific and spatially confined, pattern was recognized. A slight trend towards negative correlations was observed between the extent of CpG methylation between -10kb and -500bp upstream of the TSS. Right at the TSS, a strong negative correlation was observed. In contrast, in the downstream regions of the TSS from 500bp to 5kb a slight trend towards positive correlations can be observed. These observations are in contrast to the observations made across maize inbreds, where methylation at CpG DMRs associated with repressed expression in gene body regions [34].

The trend of association between methylation at CHG DMRs was largely consistent with that of CpG DMRs in the upstream region of the TSS from -5kb to -500bp and at the TSS from -500kb to 500kb. However, the trend of association between methylation at CHG DMRs in the downstream regions from 500bp to 5kb was strongly negative and consistent with the findings in maize [34]. This may be explained by the association of H3K9me2 with CHG methylation, though this is subject for further investigation due to conflicting studies in the past [82, 83].

Methylation at DMRs of the CHH context were positively correlated with gene expression throughout all distance groupings. This trend towards positive correlation was less prominent in the distal regions from the TSS and consistent with the observations in one barley inbred line and with previous observations in maize [34].

In conclusion, differences in gene expression are associated with more specific spatially confined differences in methylation across inbred lines than across different genes in one inbred line, especially in the CpG and CHG context.

### Association of DMR and phenotypic variation

Assessing the association of DMRs as a characteristic of genotypes with phenotypic variation is an important step to understand the origin of the missing heritability better. It is well known that *VRN-H1* is a key regulator of flowering time in cereal crops and one of the main actors in vernalization response [68, 98]. Previous research illustrated that a large deletion of around 5kb in an intronic region of *VRN-H1* in spring barley compared to winter barley is one of the reasons for an increased basal expression of *VRN-H1* in non vernalized seedlings [98]. Morex, a spring barley variety, showed high levels of histone 3 lysine 4 trimethylation (H3K4me3), a histone modification associated with increased transcription, at *VRN-H1*. Our results suggest that not only a deletion in the intronic region determines the expression of *VRN-H1* among inbred lines, but also DNA methylation. We identified within spring barley two epialleles where a increased CHG methylation lead to a significantly increased *VRN-H1* expression and therefore earlier flowering time (Figure 9c). Noteworthy, this DMR was not associated with local sequence variation. The regulation of flowering time by non-CG methylation has already been revealed in winter wheat as a consequence of vernalization treatment [99]. Here we propose that non-CG DNA methylation may regulate the basal expression of *VRN-H1* as a characteristic of spring barley genotypes contributing to earlier flowering times of inbred lines carrying epiallele 1. Previous research with the inbred lines of our study has already reported a flowering time quantitative trait loci (QTL) which confidence interval included *VRN-H1* in the progeny of parents IG31424, carrying epiallele 2, and Kharsila, carrying epiallele 1 at *VRN-H1* [100].

### Methylation as a genomic predictor

Incorporating methylation information in genomic prediction has the potential to explain more of the missing heritability, and thereby getting a wider understanding of the causes of phenotypic variation, as methylation is largely independent to sequence variation (Figure 7) and also partially independent to gene expression variation (Figure 8). The prediction ability of DNA methylation data in genomic prediction models in barley was largely dependent on the predicted trait (Figure 10). About half of the examined phenotypic traits showed noteworthy improvements by using methylation information as a predictor.

These differences are presumably caused thereby that the traits differ in the importance of polymorphisms in the coding sequence versus gene expression effects. Nevertheless, our results highlight that especially the combination of DNA sequencing SNPs and methylation is valuable for many traits and performs generally better than the combination of DNA sequencing SNPs and gene expression. These observations in barley are in line with previous studies [34, 40, 101, 102].

### Summary

The average CpG and CHG methylation level of barley was found to be high in comparison to other cereal crops, though the CHH methylation level was lower than previously reported. The genome-wide pattern of barley’s methylome were consistent to that of other crops. The low extent of linkage disequilibrium between DMRs and SNPs highlights that methylation provides a unique layer of information not accessible through sequence variation and not predictable by SNPs in the majority of cases. Although the correlation between DMRs and gene expression was mostly low to moderate, about 5.69% of all annotated genes showed strong correlations with DMRs. We observed that the direction of the associations between methylation at DMRs with gene expression of the same gene across inbred lines was much more specific and spatially confined than associations of methylation with the expression of different genes in one inbred line. This was exemplified for the association of the known MADS-box transcription factor encoding gene *VRN-H1* with gene expression and flowering time. These observations underline the importance to extend prediction models to use epigenetic variation in combination with genomic predictors for a more accurate genomic prediction as exemplified in this study. In many cases methylation was able to outperform SNPs and gene expression as a predictor for phenotypic variation depending on the trait of interest, illustrating that methylation variation is one of the factors explaining the missing heritability.

## ACKNOWLEDGEMENTS

This research was supported by the Deutsche Forschungsgemeinschaft (DFG, German Research Foundation) under Germany’s Excellence Strategy—EXC 2048/1—Project ID: 39068611 and core funding of the Julius Kühn Institute.

Computational infrastructure and support were provided by the Centre for Information and Media Technology at Heinrich Heine University Düsseldorf.

We additionally thank Federico Casale for helpful discussions regarding the relationship between DNA Methylation and meiotic recombination rates.

The authors declare no conflict of interest.

---

**Fig. S1:**
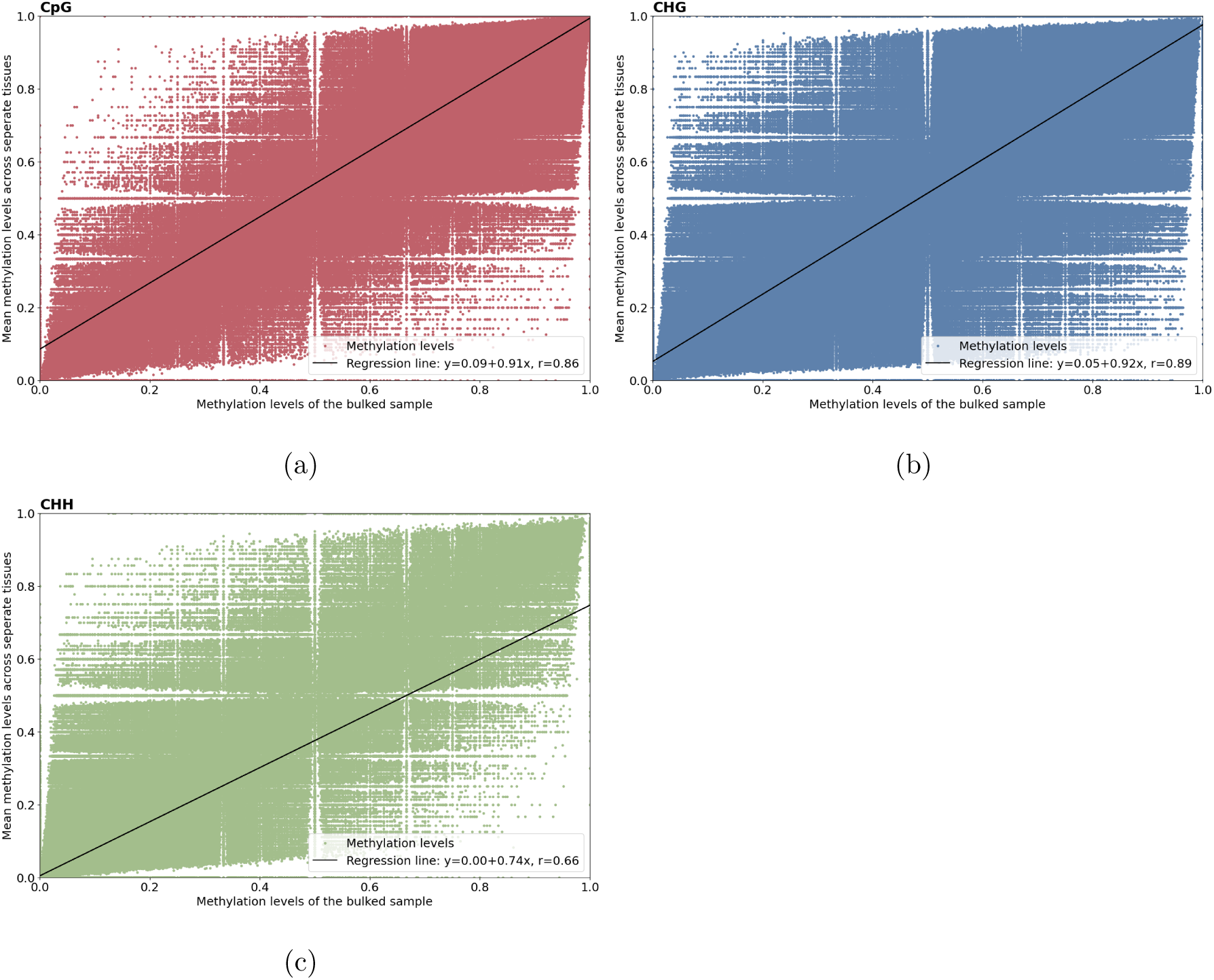
Pearson correlation of the average methylation levels across the Sissy tissue samples and the methylation levels of the bulked Sissy sample for the a) CpG, b) CHG, and c) CHH context.

**Fig. S2:**
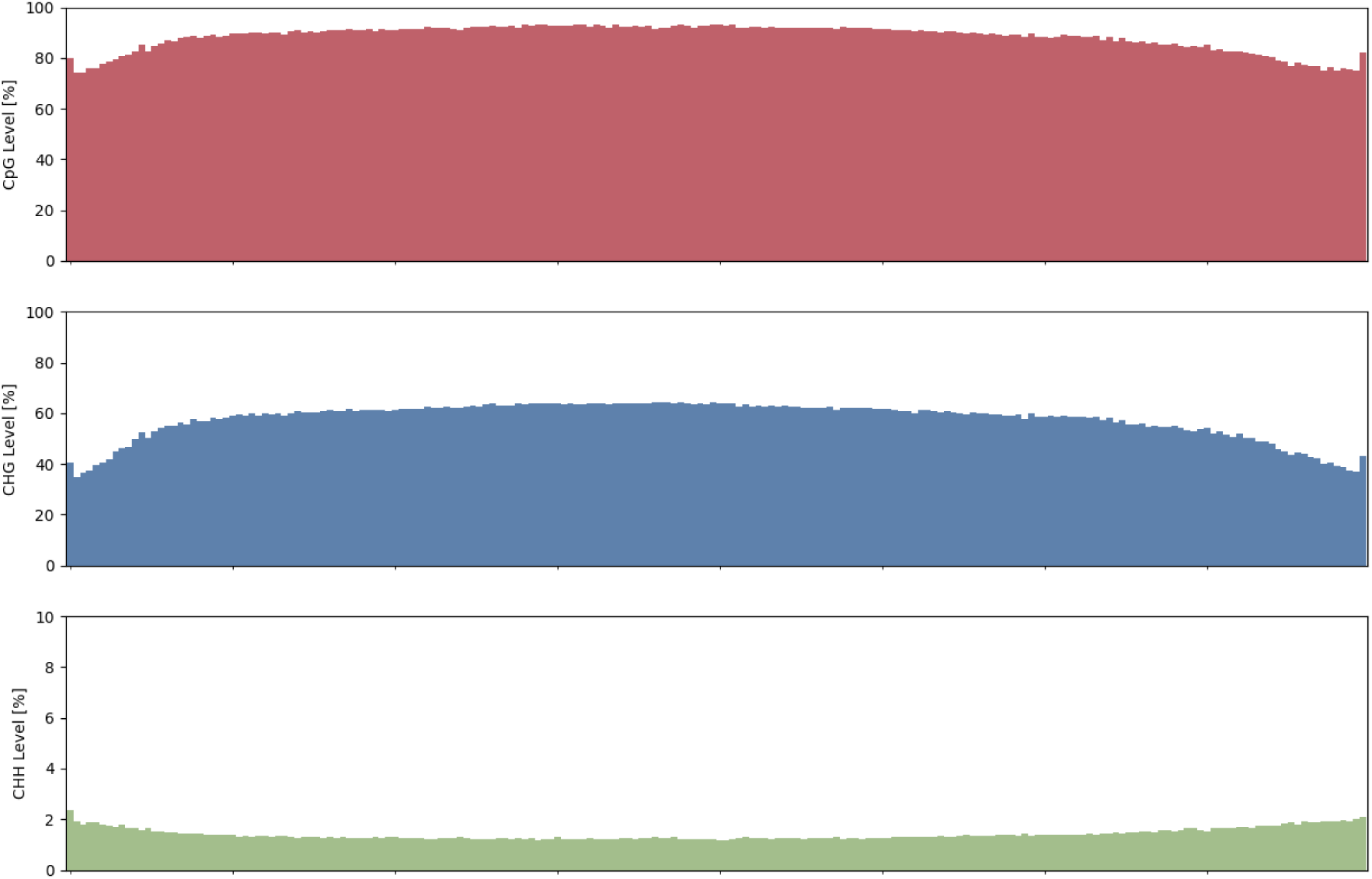
Average chromosome methylation of each sequence context calculated in 200 bins across 23 inbred lines and 7 chromosomes.

**Fig. S3:**
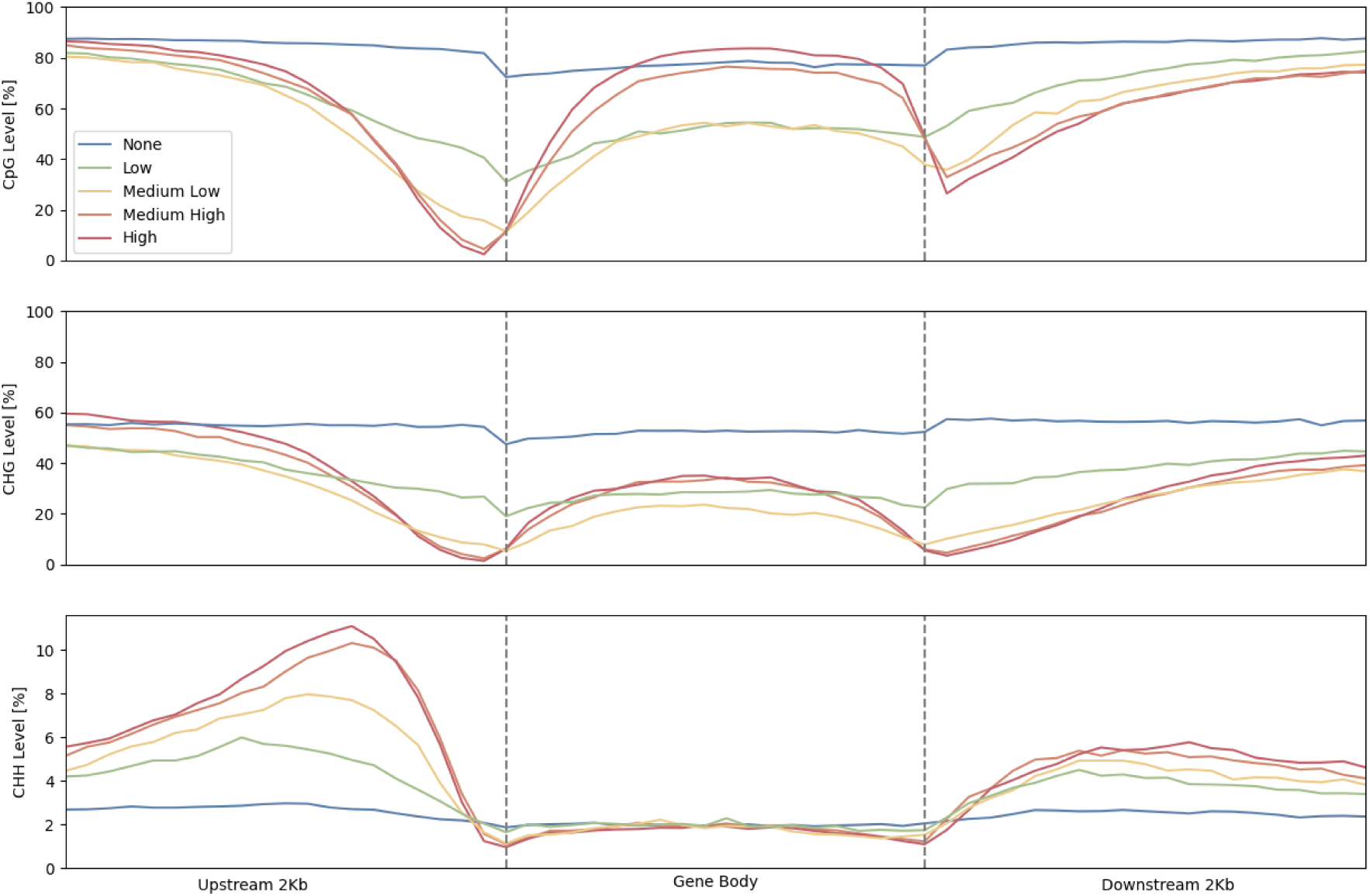
Average methylation levels of genes in Sissy’s seedling tissue categorized based on their expression levels in the same tissue (100% quantile ≥ High *>* 75% quantile; 75% quantile ≥ Medium High *>* 50% quantile; 50% quantile ≥ Medium Low *>* 25% quantile; 25% quantile ≥ Low *>* 0% quantile; None = 0).

**Fig. S4:**
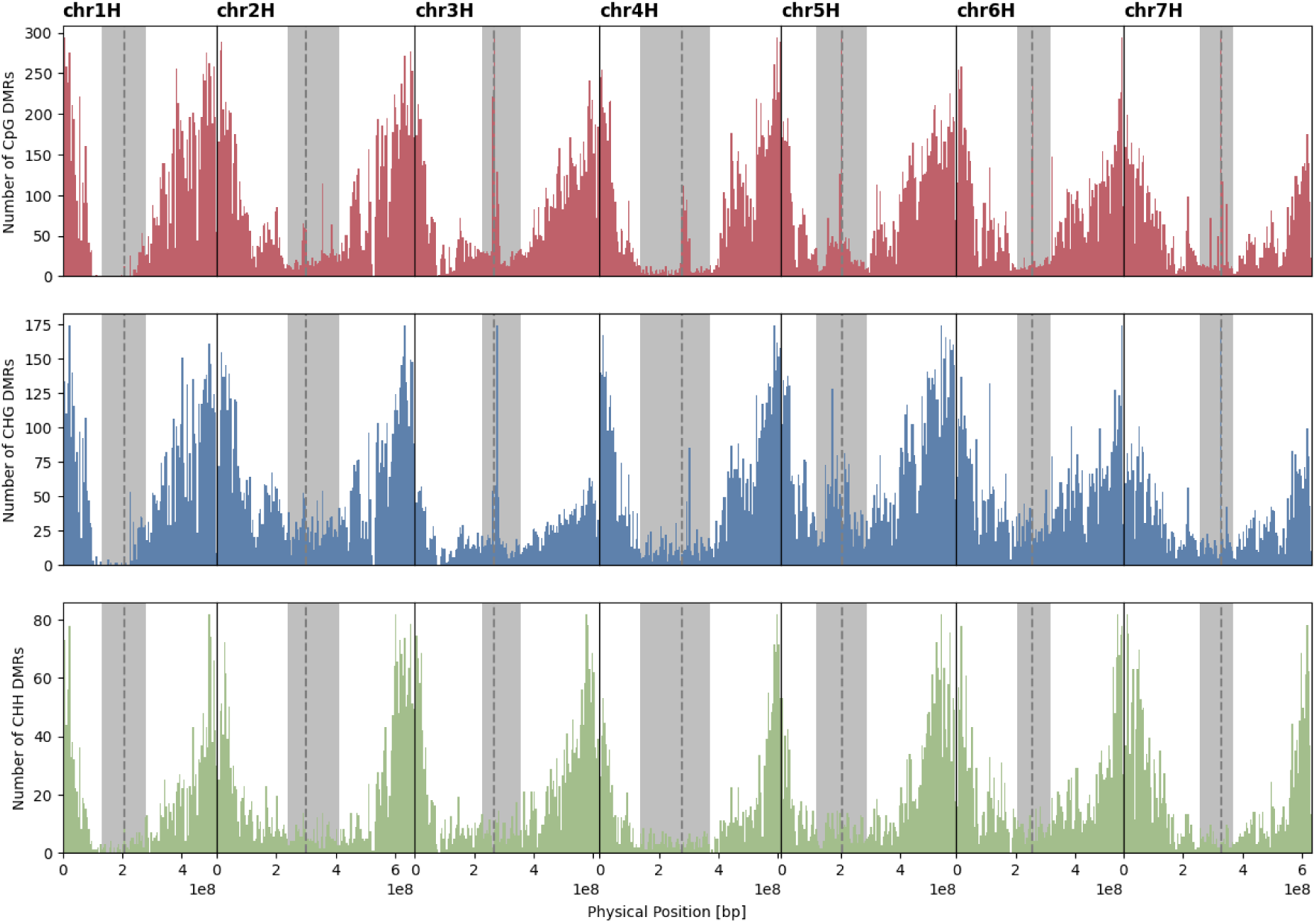
Distribution of single nucleotide polymorphism (SNP) associated differentially methylated regions (DMRs) in bins of 5 Mbp across the genome. The dashed gray lines show the middle of the centromere with the pericentromeric regions highlighted in gray.

**Fig. S5:**
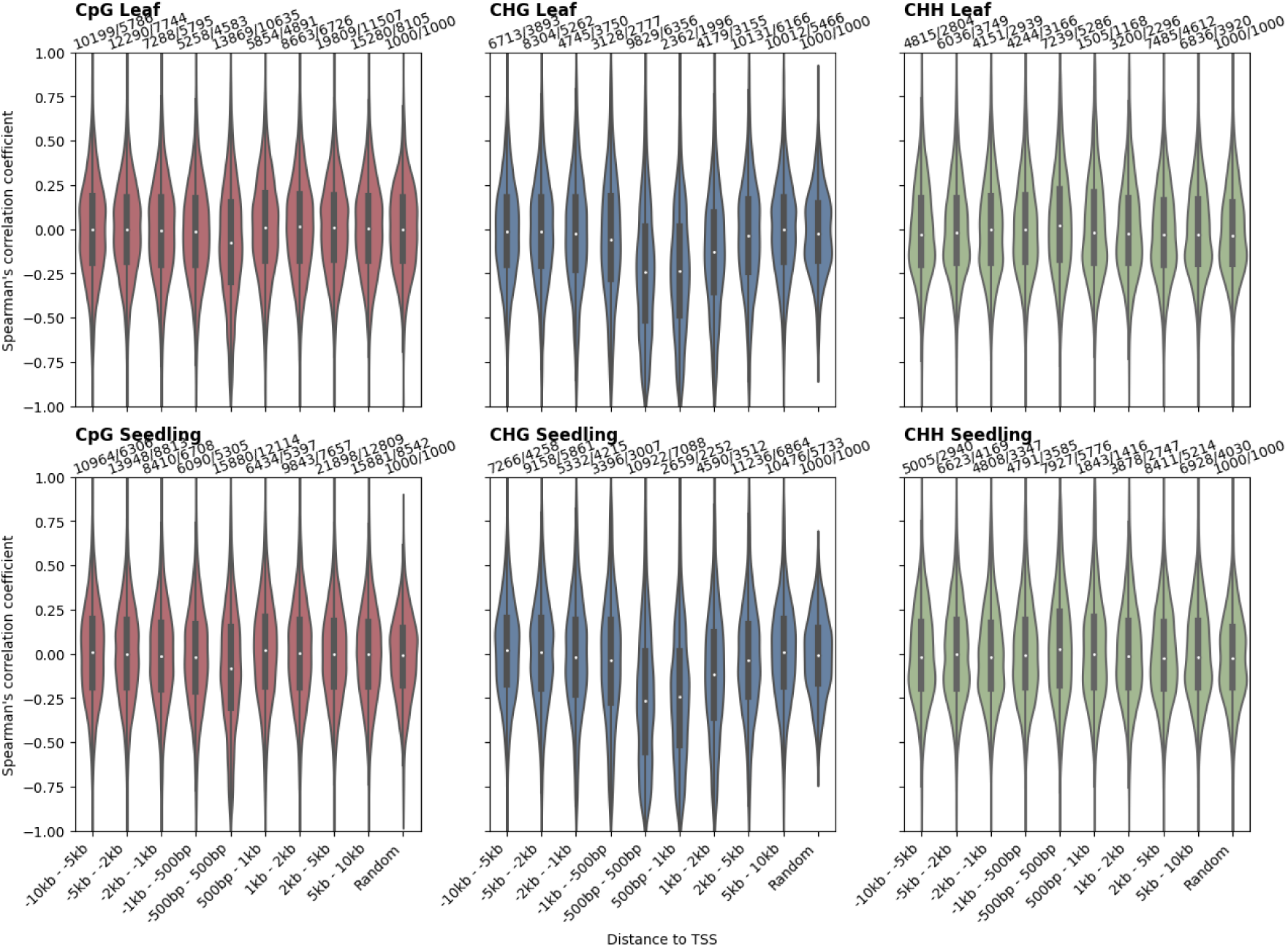
Spearman’s rank correlation coefficient distributions of differentially methylated regions (DMRs) with gene expression grouped by the DMR’s distance to the respective transcription start site (TSS) ±10 kb in intervals of 5 kb to 500 bp for the leaf and seedling expression data separately. Additionally, 1000 DMRs were randomly correlated with 1000 random genes, presented in the last category of each subplot. The numbers above the categories reflect the number of correlations, as well as the number of unique genes in the respective category.

**Table S1:**
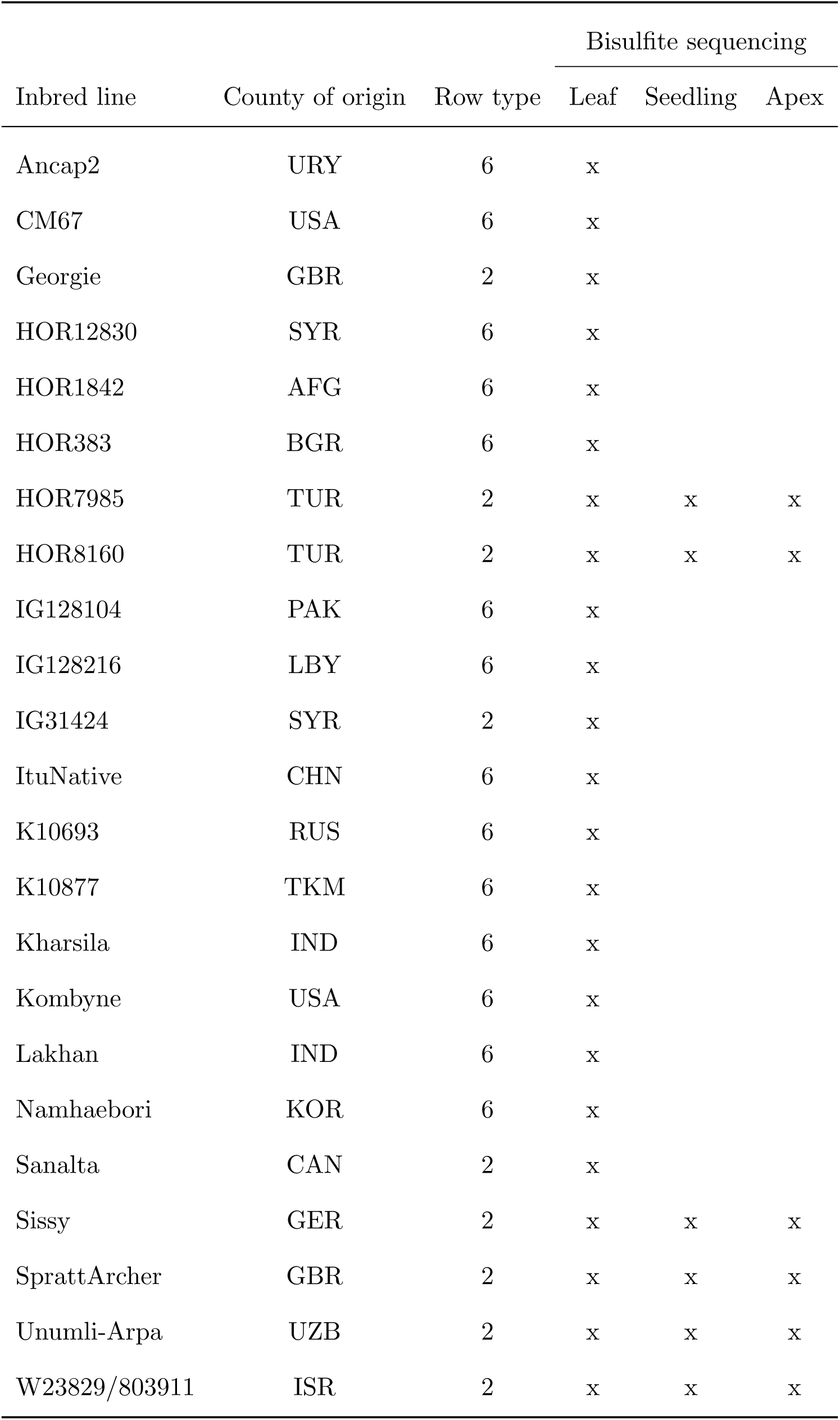
Barley inbred lines with their country of origin, row type and tissues that were part of this study.

**Table S2:** Average mapping efficiency of respective one million read subsets of the 23 barley inbreds against the uncorrected, single nucleotide polymorphism (SNP) corrected, and SNP and insertion/deletion (Indel) corrected Morex reference.

| SNP corrected | Indel Corrected | Mapping Efficiency [%] |
| --- | --- | --- |
| No | No | 59.9 |
| Yes | No | 61.8 |
| Yes | Yes | 62.2 |

**Table S3:** Average mapping efficiency of respective one million read subsets of the 23 barley inbreds mapped against the single nucleotide polymorphism (SNP) corrected Morex reference under varying parameters. Parameter values marked with an asterisk reflect default values.

| Scoring Function | Maximum Insert-size | Allowed Mismatches | Mapping Efficiency [%] |
| --- | --- | --- | --- |
| $0 + -0.2 * L^*$ | 500* | 1 | 60.0 |
| $0 + -0.2 * L^*$ | 500* | 0* | 61.8 |
| $0 + -0.2 * L^*$ | 1200 | 0* | 64.0 |
| $0 + -0.4 * L$ | 500* | 0* | 67.2 |
| $0 + -0.4 * L$ | 1200 | 0* | 69.6 |
| $0 + -0.6 * L$ | 500* | 0* | 70.0 |
| $0 + -0.6 * L$ | 1200 | 0* | 72.4 |

